# Global Analysis of Human mRNA Folding Disruptions in Synonymous Variants Demonstrates Significant Population Constraint

**DOI:** 10.1101/712679

**Authors:** Jeffrey B.S. Gaither, Grant E. Lammi, James L. Li, David M. Gordon, Harkness C. Kuck, Benjamin J. Kelly, James R. Fitch, Peter White

**Author notes:** Corresponding author Mailing address: The Institute for Genomic Medicine, Nationwide Children’s Hospital, 575 Children’s Crossroad, Columbus, OH 43215. USA. Department of Pediatrics, College of Medicine, The Ohio State University, Columbus, Ohio, USA.

## Abstract

**Background:** In most organisms the structure of an mRNA molecule is crucial in determining speed of translation, half-life, splicing propensities and final protein configuration. Synonymous variants which distort this wildtype mRNA structure may be pathogenic as a consequence. However, current clinical guidelines classify synonymous or “silent” single nucleotide variants (sSNVs) as largely benign unless a role in RNA splicing can be demonstrated.

**Results:** We developed novel software to conduct a global transcriptome study in which RNA folding statistics were computed for 469 million SNVs in 45,800 transcripts using an Apache Spark implementation of ViennaRNA in the cloud. Focusing our analysis on the subset of 17.9 million sSNVs, we discover that variants predicted to disrupt mRNA structure have lower rates of incidence in the human population. Given that the community lacks tools to evaluate the potential pathogenic impact of sSNVs, we introduce a “Structural Predictivity Index” (SPI) to quantify this constraint due to mRNA structure.

**Conclusions:** Our findings support the hypothesis that sSNVs may play a role in genetic disorders due to their effects on mRNA structure. Our RNA-folding scores provide a means of gauging the structural constraint operating on any sSNV in the human genome. Given that the majority of patients with rare or as yet to be diagnosed disease lack a molecular diagnosis, these scores have the potential to enable discovery of novel genetic etiologies. Our RNA Stability Pipeline as well as ViennaRNA structural metrics and SPI scores for all human synonymous variants can be downloaded from GitHub https://github.com/nch-igm/rna-stability.

## Introduction

While next generation sequencing (**NGS**) has accelerated the discovery of new functional variants in syndromic and rare monogenic diseases, many more disease-causing genes and novel genetic etiologies remain to be discovered [1, 2]. Accurate molecular genetic diagnosis of a rare disease is essential for patient care [3], yet today’s best molecular tests and analysis strategies leave 60-75% of patients undiagnosed [4-8]. Current clinical practice for sequence variant interpretation focuses primarily on missense, nonsense or canonical splice variants [9], with numerous bioinformatics prediction algorithms and databases developed for functional prediction and annotation of non-synonymous single-nucleotide variants (nsSNVs) that impact protein function through changes in the underlying coding sequence [10]. However, these algorithms are inadequate to infer pathogenicity in non-protein-altering variants such as intronic or synonymous variants, which are under different and weaker evolutionary constraints [11]. While the potentially pathogenic impact of non-synonymous single nucleotide variants (**nsSNVs**) that change the protein sequence are well understood, we have limited knowledge in regard to the role that synonymous SNVs (**sSNVs**) may have in human health and disease.

Synonymous variants result in codon changes that do not alter the amino acid sequence of the translated protein and as such for decades were referred to as “silent.” However, there is a growing body of evidence demonstrating that synonymous codons have vital regulatory roles [12-16] among the most important of which is their contribution to RNA structure.

Messenger RNA (**mRNA**) is a single-stranded molecule that adopts three levels of structure: the *primary sequence* forms base pairs among its own nucleotides to build the *secondary structure*, which further folds through covalent attractions to form the *tertiary structure* (**F****igure** **1**) [17]. While the tertiary structure of mRNA is challenging to model and poorly understood, sophisticated tools exist to compute the ensemble of possible secondary structures and determine the optimal structure for a given mRNA strand [18].

**FIGURE 1.**
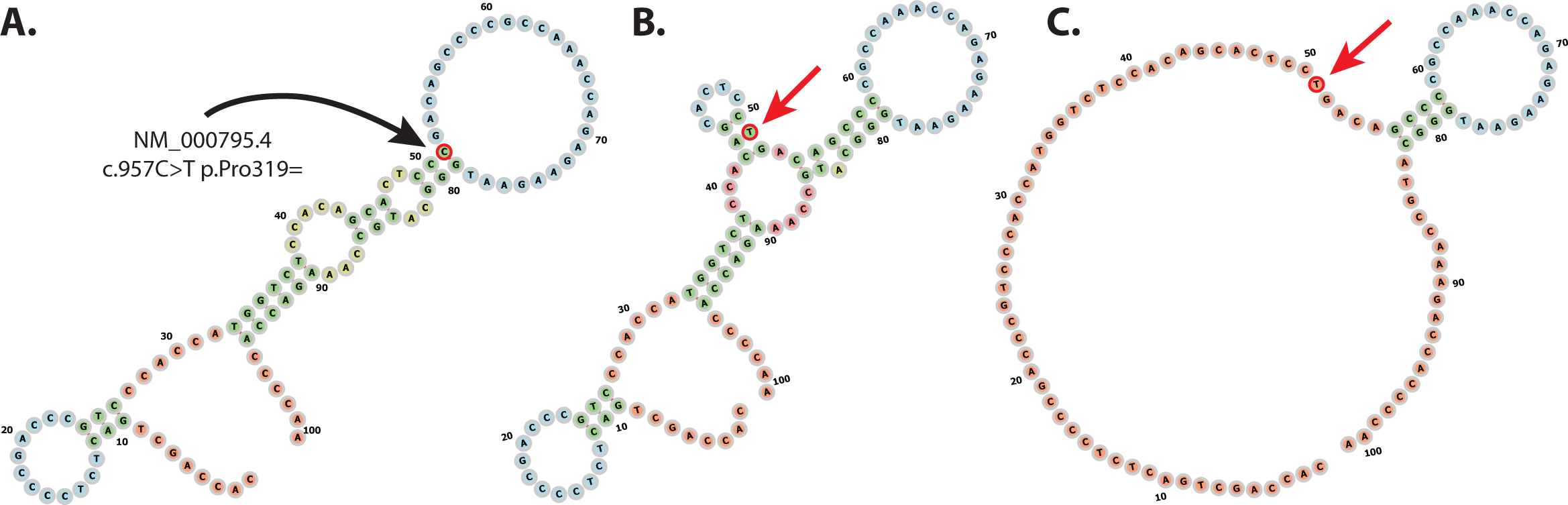
A synonymous variant introduces a marked change in local minimum free energy of the mRNA secondary structures in the *DRD2* gene. Using a known synonymous variant of pharmacogenomic significance in the dopamine receptor, DRD2 (NM_000795.4:c.957C>T (p.Pro319=)), this figure demonstrates how the 101-bp window used in our analysis captures the variant’s impact on RNA secondary structure. Wildtype (**A**) and mutant (**B** and **C**) sequences (RefSeq transcript NM_000795.4, coding positions 907-1008) are identical except for a synonymous C->T mutation at position 51 (major “C” allele is indicated by the black arrow, minor “T” allele is indicated by the red arrow). **(A)** Wildtype optimal and centroid structures (which coincide) demonstrate a relatively stable secondary structure with a minimum free energy of −12.5 kcal/mol. In the ensemble of possible structures arising from the sSNV a position 51, there is a significant reduction in stability of the molecule in terms of both the (**B)** mutant optimal structure (−11.5 kcal/mol) and (**C)** mutant centroid structure (−5.1 kcal/mol). The synonymous variant results in a less stable mRNA molecule which laboratory studies demonstrate reduces the half-life of the transcript, ultimately reducing protein expression of the dopamine receptor, DRD2. Nucleotides are colored according to the type of structure that they are in: Green: Stems (canonical helices); Red: Multiloops (junctions); Yellow: Internal Loops; Blue: Hairpin loops; Orange: 5’ and 3’ unpaired region.

Studies first published in 1999 indicated that stable mRNA secondary structures are often selected for in key genomic regions across all kingdoms of life [19-22]. Synonymous variants impacting RNA structure can alter global RNA stability, where stable mRNAs tend to have longer half-lives and less stable RNA molecules may be more rapidly degraded resulting in lower protein levels [23-28]. The stability of an mRNA transcript affects translational initiation and can determine how quickly a given protein is translated [19-21, 29-31]. Recent studies strongly linked mRNA structure to protein confirmation and function, with synonymous codons acting as a subliminal code for the protein folding process [16, 30, 32-35]. mRNA structure can also facilitate or prevent miRNAs and RNA-binding proteins from attaching to specific structural motifs [36-39]. Given these multiple mechanisms, when synonymous variants are ignored, we are almost certainly missing novel plausible explanations for genetic disease.

The role of mRNA structure in human health and disease, however, is poorly comprehended and relatively few pathogenic variants impacting mRNA folding have been described [23, 24, 26, 28]. A structure-altering sSNV in the dopamine receptor DRD2 was shown to inhibit protein synthesis and accelerate mRNA degradation [40]. A sSNV in the *COMT* gene, implicated in cognitive impairment and pain sensitivity, was shown *in vitro* to constrain enzymatic activity and protein expression [41]. A sSNV discovered in the OPTC gene of a glaucoma patient resulted in decreased protein expression *in vivo* [42]. In cystic fibrosis patients, a sSNV in the *CFTR* gene was linked to decreased gene expression [43]. Additionally, a silent codon change, I507-ATC→ATT, contributes to CFTR dysfunction by a change in mRNA secondary structure that alters the dynamics of translation leading to misfolding of the CFTR protein [25, 27]. Two sSNVs in the NKX2-5 gene, identified in patients with congenital heart disease, decreased the mRNA’s transactivation potential in a yeast-based assay [44]. In hemophilia B, the sSNV c.459G>A in factor IX impacts the transcript’s secondary structure and reduces extracellular protein levels [45], and both synonymous and nonsynonymous SNVs were shown more likely be deleterious when occurring in a stable region of mRNA in hemophilia associated genes F8 and Duchenne’s Muscular Dystrophy [46].

We hypothesize that these reported instances of mRNA structure playing a role in disease represent only the tip of the iceberg and that many undiagnosed genetic disorders might also be influenced by disruptions to mRNA structures. As such, the goals of this study were the creation of metrics to predict a sSNV’s pathogenicity due to its effects on mRNA structure and to utilize these metrics to test the hypothesis that synonymous variants predicted to have disruptive impacts on RNA stability would show significant constraint in the human population. In successfully doing so we hope to provide the genetics research community with tools to identify novel genetic etiologies in both monogenic genetic disorders and more complex human disease, thus leading to improved diagnosis and the possibility of novel prevention and treatment approaches.

## Results

### Massively parallel generation of RNA stability metrics

Global assessment of sSNVs is truly a big data problem as it requires generation and evaluation of several raw values for each of hundreds of millions of positions within the genome. To address this challenge and successfully predict the mRNA-structural effects of every possible sSNV, we developed novel software built upon the Apache Spark framework (**F****igure** **2**). Apache Spark is a distributed, open source compute engine that drastically reduces the bottleneck of disk I/O by processing its data in memory whenever possible [47]. This leads to a 100x increase in speed and allows for more flexible software design than can be achieved in the traditional Hadoop MapReduce paradigm. Spark is well suited to address many of the challenges faced in analyzing big genomics data in a highly scalable manner and adoption is growing steadily, with applications such as SparkSeq [48] for general processing, SparkBWA [49] for alignment and VariantSpark for variant clustering [50]. By developing a solution within this framework, we eliminate significant computational hurdles standing in the way of large-scale analysis of sSNVs.

**FIGURE 2.**
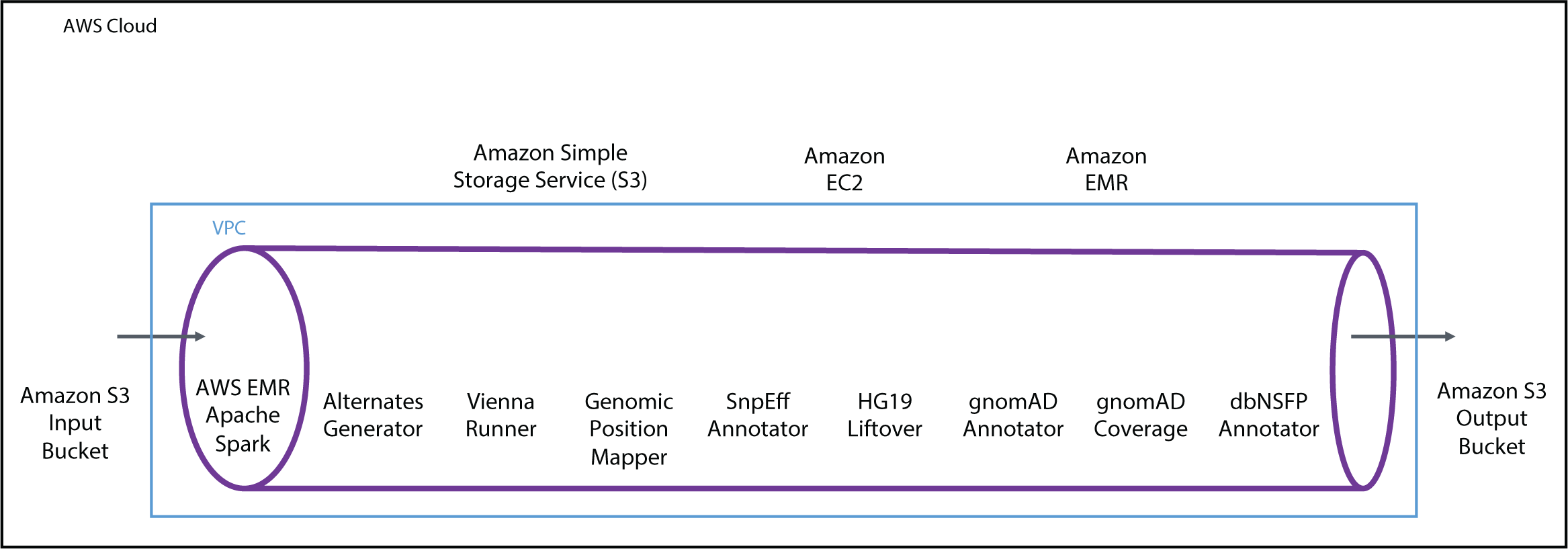
Graphical depiction of computational workflow used to generate ViennaRNA folding metrics for the entire transcriptome. The entire analysis workflow was parallelized using Apache Spark and the Amazon Elastic Map Reduce (EMR) service, generating 5 billion ViennaRNA metrics over the course of 2 days. Using a custom pipeline developed for the process that was executed across 47 Amazon Elastic Cloud Compute (EC2) spot instances, input data was retrieved from an Amazon Simple Storage Solution (S3) bucket and processed through the pipeline consisting of 8 steps. We first obtained the 101-base sequence centered around a SNV in a transcript and generated three alternate sequences (with the ALT rather than the REF at position 51) (step 1). We next applied ViennaRNA modules to sequence to obtain structural metrics (step 2). Results were then mapped to chromosomal coordinates (step 3) and annotated with SnpEff to identify splice variants (step 4), lifted to the hg19 build (step 5), annotated with gnomAD population frequencies (step 6) and coverage information (step 7), and finally annotated with metrics from dbNSFP (step 8). Final dataset was written to Amazon S3 in Parquet columnar file format for further analysis and interpretation.

We used the RefSeq database (Release 81, GRCh38) as the source for all known human coding transcript sequences. At each position within a given transcript, four 101-base sequence windows were built, differing only in their central nucleotide, which was set to the reference nucleotide or one of the three possible alternate bases. Using Apache Spark in the Amazon Web Services (**AWS**) Elastic Map Reduce (**EMR**) service, we developed a massively parallel implementation of the ViennaRNA Package to analyze the four possible sequences. This enabled us to examine changes in mRNA folding that result from any given polymorphism, and thereby obtain ten metrics which quantified the SNV’s effect on mRNA secondary structure (see **Supplementary Table 1**). First, we utilized RNAfold to obtain predicted free energies for both mutant and wildtype sequences, which we compared directly to obtain four metrics describing the sSNV’s effect on mRNA stability. Next, we fed the predicted structures from RNAfold into the ViennaRNA programs RNApdist and RNAdistance to obtain 6 additional metrics quantifying the change in base-pairing and ensemble diversity due to each SNV. We performed this procedure for all 469 million possible SNVs in 45,800 transcripts.

After pre-processing we assigned each SNV a classification based on the most deleterious role it played in any transcript, in decreasing order of deleteriousness: start loss, stop gain, start gain, stop loss, missense, synonymous, 5 prime UTR, 3 prime UTR. We then focused on the set of 22.9 million synonymous variants. While non-synonymous variants also play a role in mRNA structure, we chose to exclude 63.8 million nsSNVs from the subsequent analysis as their impact on conserved amino acid sequences would make it difficult to discern constraint at the mRNA structural level. We also filtered out variants implicated in splicing or lacking annotations needed in future steps, leaving us with a core dataset of 17.9 million sSNVs (see **Methods** for details, **F****igure** **2** for a summary of our computational pipeline, and **Supplementary Table 2** for a record of the number of SNVs filtered at each stage). Of the 10 mRNA-structural metrics computed for each sSNV we adopted three as the primary focus for our analysis: dMFE, CFEED, and dCD. The metric dMFE (delta Minimum Free Energy) measures the change in overall mRNA stability imputed by the sSNV, while CFEED (Centroid Free Energy Edge Distance) gives the number of base pairs that vary between the mutant and wildtype centroid structures. The metric dCD (delta Centroid Distance) measures the sSNV’s effect on the diversity of the mRNA’s structural ensemble.

To test whether certain sSNVs are under constraint due to their effect on mRNA structure, RNA folding metrics from our ViennaRNA pipeline were combined with population frequencies from the Genome Aggregation Database (gnomAD), containing aggregate WGS and WES data from a total of 138,632 unrelated human individuals [51]. Our expectation was that SNVs with disruptive structural properties would be found less frequently in human population. Constrained variants were defined as those absent from gnomAD, versus un-constrained variants as being those with exon minor allele frequency (**MAF**) > 0, a strategy similar to that employed by other groups [52, 53].

### Global constraint to maintain stability

Our study reveals a striking connection between a given SNV’s impact on mRNA structure and its frequency in the gnomAD database. We define the central variable Y to be Y=1 when a SNV is present in gnomAD and Y=0 when the SNV is absent. We hypothesize that variants which disrupt mRNA structure should have Y=0 (i.e. be absent from the gnomAD database), while those with limited impact on structure should tend to have Y=1 (i.e. appear at least once in the gnomAD database).

This central hypothesis is validated in **F****igure** **3**, which depicts the proportion of SNVs with Y=1 at every value of our stability-metric dMFE. All four variant classifications – synonymous, 5 prime UTR, 3 prime UTR and missense – show a bi-directional constraint to maintain the wild-type mRNA structure. In each context the bell-shaped pattern indicates that a SNV is most likely to appear in the population when it maintains the existing level of stability i.e. has dMFE close to 0. When the SNV either over-stabilizes the mRNA (low dMFE) or de-stabilizes it (high dMFE) the SNV is depleted in the population roughly in proportion to the level of disruption. While this pattern of constraint was observed across all four variant classes, **F****igure** **3** indicates that synonymous variants experience the most extreme constraint for mRNA structure. This justifies our decision to regard the synonymous case as prototypical, and henceforth we restrict our attention to it.

**FIGURE 3.**
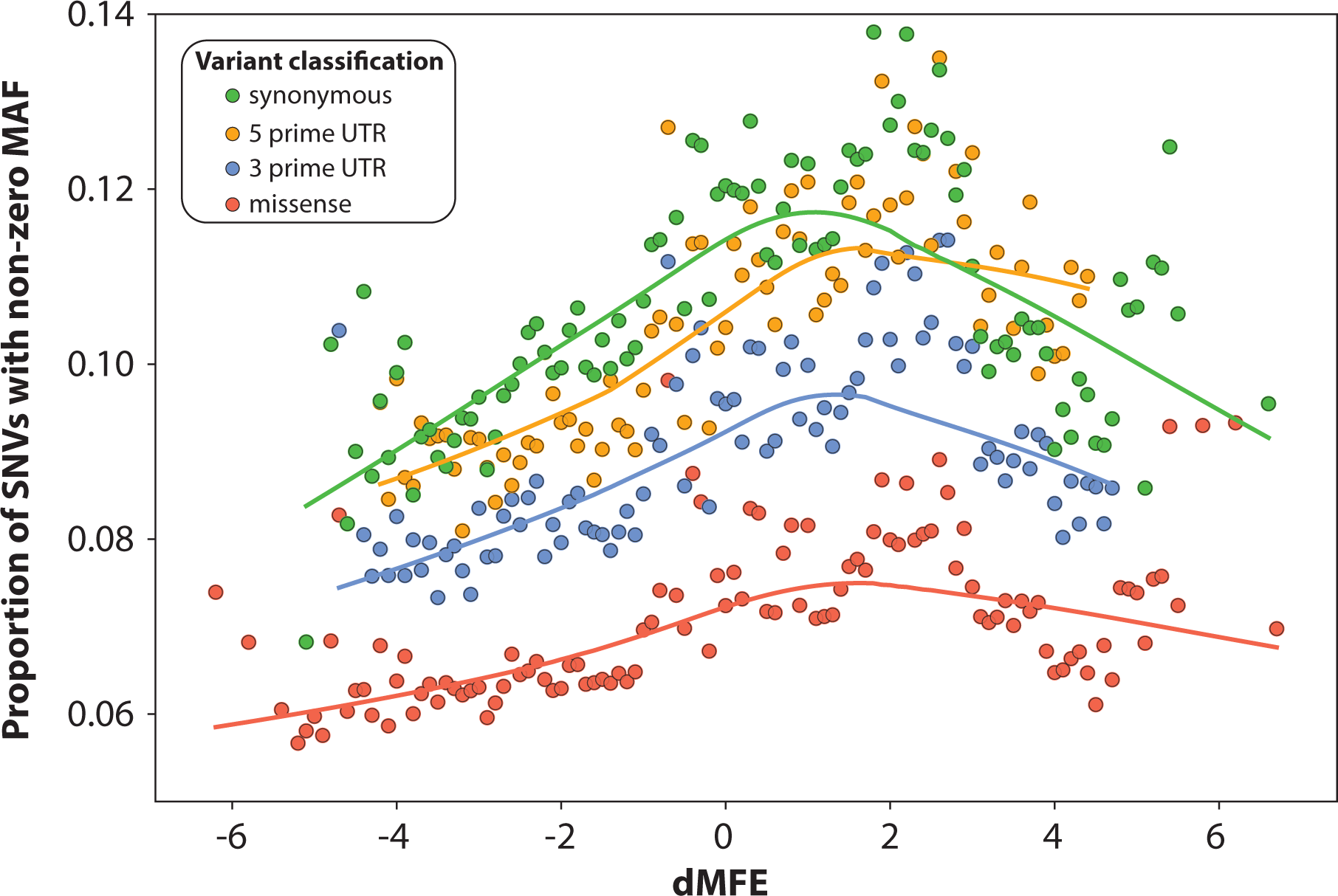
Exonic SNVs predicted to impact mRNA structure are constrained in the human population. Population frequency of SNVs was plotted against predicted impact on mRNA structure. Circles show proportion of SNVs with nonzero gnomAD exonic frequency at each value of the RNA stability metric dMFE. The bell-shaped pattern of constraint was observed across all classes of SNVs, with constraint appearing to be greatest in sSNVs (green), followed by SNVs in the 5 prime UTR (orange), then SNVs in the 3 prime UTR (blue), and finally nsSNVs (red). Values of dMFE with fewer than 1000 (synonymous), 200 (UTRs) or 2000 (missense) positive-MAF sSNVs are excluded. Axis limits are restricted to show main pattern more clearly, resulting in removal of a few high-P(MAF>0) synonymous outliers. Only SNVs with high exonic coverage are represented (see **Methods** for details.)

**F****igure** **4** depicts the effect of mRNA-structural changes for sSNVs. **F****igure** **4A** (also depicted by green circles in **F****igure** **3**) shows the correlation between Y and the stability metric dMFE across all possible sSNVs. The bell-shaped distribution shows that any change to mRNA stability bidirectionally decreases the chance of the SNV’s appearing in a human population.

**FIGURE 4.**
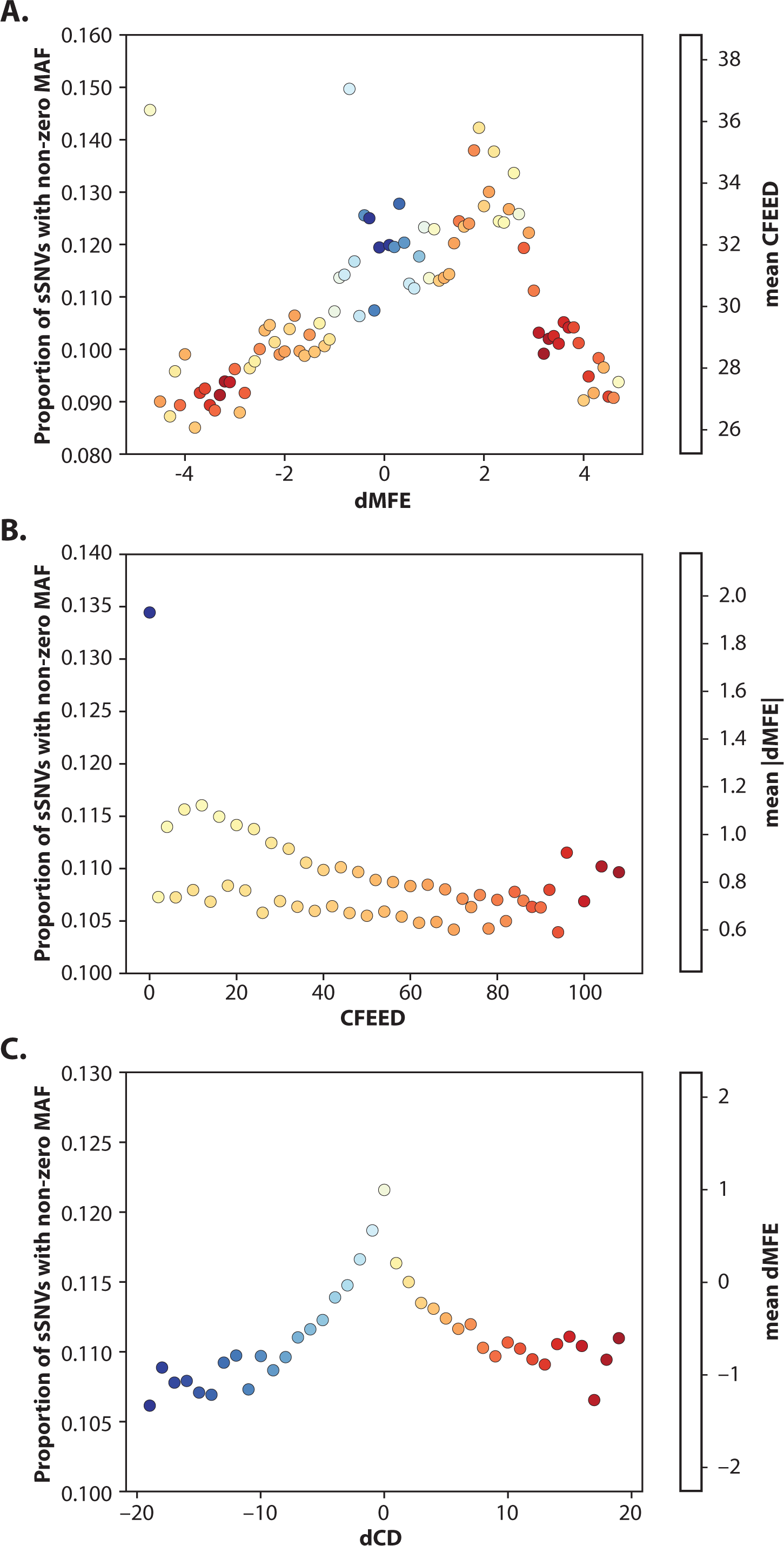
Synonymous variants predicted to impact mRNA structure are constrained in the human population. Population frequency of sSNVs were plotted against the predicted impact on mRNA structure. Synonymous variants that disrupt structure tend to be absent from the gnomAD database, while those with limited impact on structure appear at least once in the gnomAD database. **(A)** Proportion of sSNVs with nonzero gnomAD frequency at each value of the RNA stability metric dMFE. Points with fewer than 2000 positive-MAF sSNVs excluded. Color represents average CFEED value, to highlight the relationship between minimum free energy and edit distance. **(B)** Analogous plot for metric CFEED measuring edge differences between mutant/wildtype centroid structures. Color represents |dMFE|, measuring absolute change in stability. **(C)** Analogous plot for diversity-metric dCD measuring change in structural ensemble diversity due to sSNV. Color is by dMFE measuring change in stability.

**F****igure** **4B** shows an analogous plot for the structural disruption metric CFEED (see **Supplementary FIGURE 2** for an illustration of how CFEED is calculated). This plot appears to depict two separate trends, but actually shows a single pattern that alternates between high and low on successive values: the sSNVs with CFEED=0,4,8,12… are enriched over those with CFEED=2,6,10,14… (CFEED can only take on even values because the destruction/creation of a base pair always requires two edits). One possible explanation for this duality is that when CFEED fails to be divisible by 4, there is necessarily a change in the total number of base-pairings in the mRNA centroid structure. Thus, sSNVs which conserve the total number of base-pairs could be potentially favored. **F****igure** **4B** also supports the hypothesis that structurally disruptive sSNVs should appear less frequently in the population. We see that sSNVs which leave the centroid structure unchanged (i.e. CFEED=0) are roughly 15% more common than those sSNVs predicted to alter it. And within each of the two separate trends (that is, the multiples and non-multiples of 4) the population frequency declines as the number of centroid base-pairing changes grows from small to large.

Finally, sSNVs which either diversify the ensemble of mRNA structures (high dCD) or homogenize it (low dCD) are depleted in the population proportionately to their disruptions, as shown in **FIGURE 4C**. The symmetry in depletion between over- and under-diversifying sSNVs is surprisingly regular. (Here and throughout, dCD values are rounded to the nearest integer, a step necessitated by the fact that the dCD metric does not cluster into just a few values like the other two.)

The relationship between the three metrics is illuminated by color-coding in **F****igure** **4.** We observe in **F****igures** **4A** **AND** **4B** that disruptions in the magnitude of stability (|dMFE|) and base-pairing (CFEED) of a sSNV are markedly correlated, with the two metrics enriched for each other at extreme values (red coloring). **F****igure** **4C** depicts a clear relationship between diversity and stability, with those sSNVs diversifying the ensemble (high dCD) also tending to de-stabilize it (red). This diversity-instability relationship is intuitive, as a destabilizing mutation “frees up” portions of the mRNA to assume new forms. Together, these observations validate the central hypothesis that sSNVs which disrupt mRNA structure should be constrained in human populations.

### Variation of constraint with REF>ALT context

An mRNA’s secondary structure is largely determined by its primary structure (i.e. by the sequence of nucleotides). We therefore expect the constraint in **FIGURE 4** to be partially mediated by the sequence features around each sSNV, the most immediate of which are its reference and alternate bases. Accordingly we divide our sSNVs into 14 classes (**Table 1):** 12 classes based on their reference and alternate alleles (e.g. A>C, C>G, T>C, etc.) and 2 additional classes based on potential loss of methylated cytosine (CpG>TpG or CpG>CpA, the latter of which results from a deamination on an antisense strand). Then within each REF>ALT context we reconstruct the three plots of **FIGURE 4** and also perform weighted linear and quadratic regressions between the three different stability metrics and Y=1 (see **METHODS** for details). All significant results (p < 0.005) of this procedure appear in **Table 1**.

Looking at **Table 1A** (which shows the results for dMFE) we find that disruptions to mRNA stability are constrained across many of our sSNV classes. The fact that most of linear p-values are much smaller than the quadratic p-values indicates that in most contexts the dMFE-Y relationship is *linear*, in contrast to the bell-shaped relationship we see when considering global dMFE (**F****igure** **4A**). Therefore, the slope of the regression line indicates which direction of dMFE is enriched for Y=1. For example, in the context of G>T the negative normalized slope indicates that lower dMFE values (i.e. stabilizing) are less constrained (i.e. Y=1). The slope of the regression line (and the relationships it models) proves to depend largely on whether a context’s REF and ALT nucleotides are “strong” (C,G) or “weak” (A,T) binders. We note from **Table 1A** that strong>weak mutations consistently have negative slopes (except in the irregular context G>A; see *Constraint for mRNA stability in non-CpG-transitional contexts*), while the two weak>strong contexts A>G and T>C have positive slopes.

In **Table 1B** we observe the constraint for the structural disruption metric CFEED. The results here are surprising – the contexts are split between positive and negative slopes. In support of our hypothesis, four of the sequence contexts display a negative slope, implying that sSNVs with high CFEED values are constrained. However, in contrast to our hypothesis, three of the sequence contexts have a positive slope, which implies that sSNVs in these contexts with high CFEED values are enriched. In the case of CpG>TpG mutations the low quadratic p-value indicates that the pattern is actually bell-shaped, with both low and high CFEED values being depleted; but in G>A and C>A contexts the quadratic term is not significant. Actual plots of these patterns reveal that the ones in which CFEED is depleted are more striking (see **F****igure** **5** and **Supplementary FIGURES 4-5** for plots of Y vs. CFEED in all stability-significant contexts), but this unexpected result must still be addressed. We speak more on this topic in the **DISCUSSION.**

**FIGURE 5.**
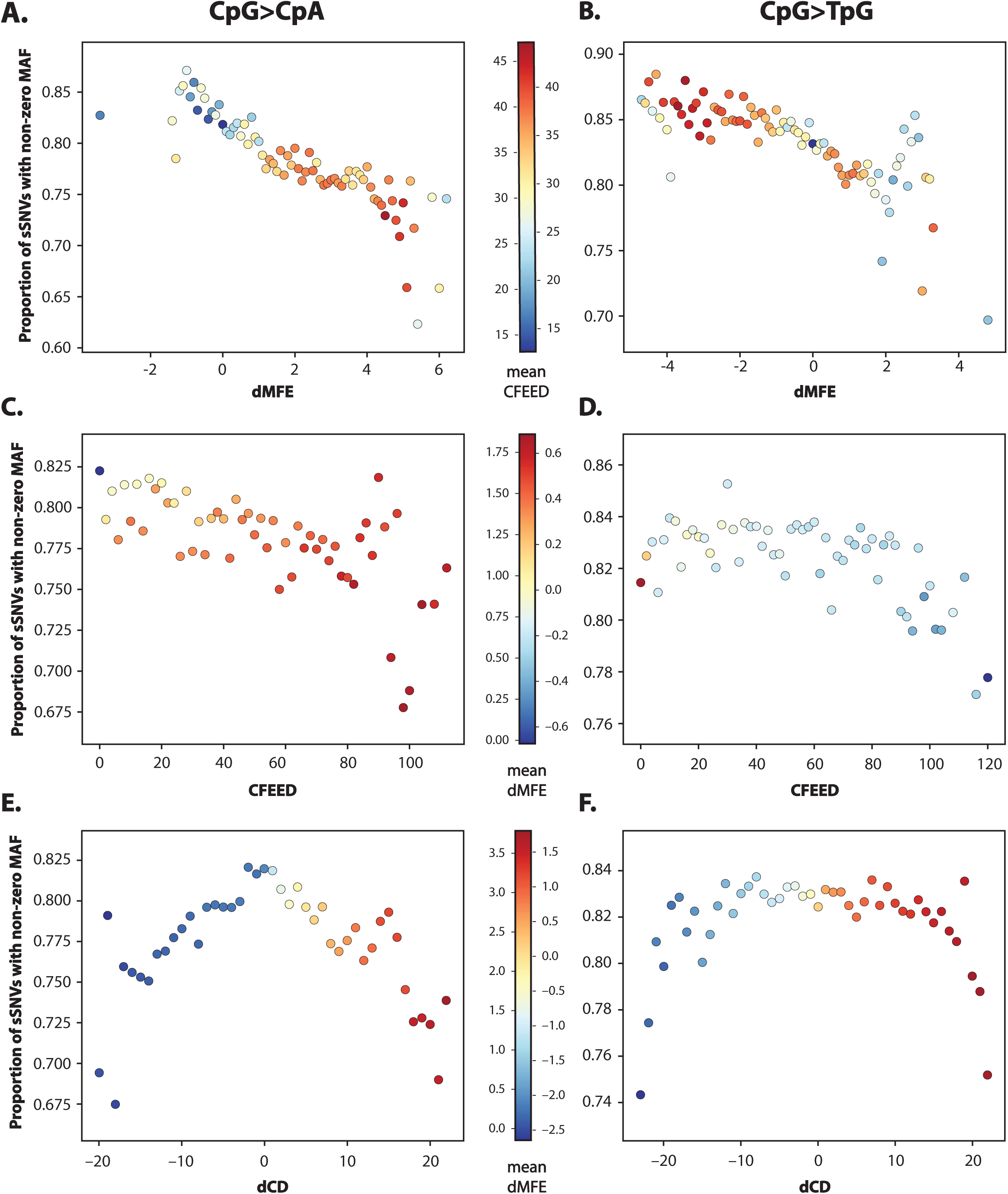
Synonymous CpG transitions are markedly constrained against destabilization of their mRNA structures. Population frequency of sSNV vs. effect on mRNA structure in synonymous CpG transitions was examined. Proportion of synonymous CpG transitions with nonzero MAF at each value of dMFE were determined for (**A**) CpG>CpA and (**B**) CpG>TpG synonymous mutations. dMFE values with fewer than 75 nonzero-MAF sSNVs are excluded. Color gives average CFEED in each context, ranging from 15 (blue) to 50 (red). Similarly, proportion of synonymous CpG transitions with nonzero MAF at each value of CFEED were determined for (**C**) CpG>CpA sSNVs and (**D**) CpG>TpG sSNVs. Color represents average dMFE and ranges from −0.8 (blue) to 1.85 (red). CFEED values with fewer than 75 nonzero-MAF sSNVs are excluded). Finally, proportion of synonymous CpG transitions with nonzero MAF at each value of dCD (after rounding to nearest integer) were determined for (**E**) CpG>CpA and (**F**) CpG>TpG sSNVs sSNVS. Color represents average dMFE and ranges from −3 (blue) to 4 (red). Rounded dCD values with fewer than 75 nonzero-MAF sSNVs are excluded.

Finally, **Table 1C** shows mutation contexts that are significantly constrained against changes to ensemble diversity. We see that only a few contexts experience this constraint. But when significant, the constraint for diversity appears to inherit the bidirectionality of **F****igure** **4C** (with the quadratic term being the most significant and the linear fit being very poor). In these contexts, decreases and increases to ensemble diversity appear to be equally harmful.

### CpG transitions have constraint against de-stabilization of their mRNA structures

The data in **Table 1** highlight that our observed constraint for mRNA structure is the greatest when considering CpG transitions. Since these variants (and their suppression) are crucial to the story of mRNA stability, it is important to have an appreciation of their role in a biochemical context. The dinucleotide CG (usually denoted CpG to distinguish this linear sequence from the CG base-pairing of cytosine and guanine) is capable of becoming methylated and then mutating by a process called “deamination” into a TG dinucleotide. While studies have demonstrated that methylated CpG residues are up to 40X times more likely to be deaminated than their unmethylated counterparts [54], mechanisms exist to enzymatically repair CpG deaminations [55, 56]. In mammals 70-80% of CpGs are methylated, which makes a CpG transition almost 5x more common than any other mutation-type among mammals (see **Supplementary DATA Table 3**) [57]. Possible explanations for the distribution and retention of CpGs in mammals have been extensively debated, with some arguing that there is no pattern of selective constraint on CpGs when compared with other dinucleotides within CpG islands [58].

The nucleotides C and G also form the foundation of mRNA secondary structures. Most of the energy of an mRNA structure lies in its “stacks” of nucleotides with the average energy of a C-G pair in a stack around 65% stronger than that of any other base-pairing [59]. Moreover, the self-complementarity of CpGs means that upstream and downstream instances can bind together and form a four-base stack which other base-pairs can then build around.

In the present study we find strong evidence that CpG transitions are constrained against de-stabilization of their mRNA structures. This striking trend is largely explained (in a statistical sense) by CpG content, i.e. number of CpG dinucleotides in the surrounding 120 nucleotides of the mRNA transcript (see “Depleter” R^2^ in **Table 1**). We distinguish CpG>CpA versus CpG>TpG transitions (the former of these usually results from a CpG>TpG deamination on an anti-sense DNA strand), as these two mutation-types show a qualitatively different constraint for mRNA structure. **F****igure** **5** shows the performance of our three main metrics in CpG-transitional contexts. Most strikingly, we find that synonymous CpG>CpA and CpG>TpG mutations both show a steady constraint against de-stabilization (high dMFE) (**F****IGURES** **5A & 5B**). Fascinatingly, both contexts exhibit a cluster of outliers in the most destructive (i.e. most de-stabilizing region), suggestive of extreme constraint borne of significant structural disruption. Though the two plots exhibit the same basic shape, the context CpG>CpA of **F****igure** **5A** shows higher de-stabilizing tendencies (higher dMFE values) and also a stronger constraint (lower P(Y=1)).

The behavior of the edge metric CFEED in these contexts is less clear-cut. In **F****igure** **5**C we see a clear pattern of constraint against mutations with high CFEED values; and the red coloring shows that such changes are, on average, de-stabilizing. But the constraint in the context CpG>TpG (**F****igure** **5D**) is much less forceful (in fact, its quadratic p-value is much smaller than its linear) and the blue coloring by dMFE shows such mutations are on average neutral or even de-stabilizing. Finally, **F****igures** **5E & 5F** show that the basic pattern of constraint for diversity in **F****igure** **4C** is reproduced and is essentially unchanged for both types of CpG transition. The coloring again indicates that mutations CpG>CpA are much more destabilizing than their CpG>TpG counterparts.

The markedly greater constraint and tendency towards de-stabilization among CpG>CpA transitions suggests they are under different selective pressures than CpG>TpG transitions, despite being largely produced by the same biochemical mechanism (a CpG>TpG deamination on either a positive- or negative-sense strand – see **Supplementary DATA Table 3**). We speculate on this disparity in the **Discussion.**

### Constraint for mRNA stability in non-CpG-transitional contexts

We see the strongest constraint for mRNA structure in CpG transitions, but we observe an analogous pattern in most REF>ALT contexts (as indicated by **Table 1**). We can classify these remaining contexts based on whether their slopes in **Table 1A** are positive or negative. **Supplementary Figure 4** shows plots of contexts where dMFE and the gnomAD variable Y are negatively correlated. In such contexts the data are consistent with the hypothesis that sSNVs which de-stabilize mRNA are constrained. Notably, all these contexts are strong>weak (or strong>strong in the case of C>G), consistent with the principle that one purpose of such nucleotides is to maintain stability. The coloring by CFEED indicates that a change in either direction is likely to alter the mRNA secondary structures.

In **Supplementary Figure 5** we show the contexts where dMFE and Y vary positively, which amounts to the claim that stabilizing mutations are constrained in these contexts. Correspondingly, we note that two out of three of these contexts are weak>strong (and the third is the unusual context G>A where SNPs that alter stability or diversity are actually enriched). The context T>C exhibits a notable constraint in either direction, an anomaly which we speculate on in the **DISCUSSION.**

### Depleter variables

In **Table 1** we provide a “Depleter” for the connection between our RNA folding metrics and gnomAD frequencies for each mutational context. The name “Depleter” signifies that each such variable is chosen so as to correlate negatively with gnomAD (which is why these variables are given with +/- signs in **Table 1**). For example, the Depleter for dMFE in the context CpG>CpA is +CpG content, meaning that when CpG content increases in this context, the varaible Y is depleted.

The Depleter is chosen to be the variable that best explains the connection between the mRNA structural variable and Y in the given context. The proportion of connection explained is given by the field “Depleter R^2^”. For example, in the context CpG>CpA we can explain 78% of the dMFE-gnomAD connection using a model that relies only CpG content.

To determine which variable is most informative (and should therefore be called the Depleter) we compute an associated R^2^ for a set of features of the sequence around the sSNV (the upstream/downstream nucleotides and the proportion of A, C, G, T, CpG or ApT [di]nucleotides in the surrounding 120 bases). Each of these features is used to build a simple logistic model to predict Y=1, and the predictions of the model are then compared to the actual proportion P(Y=1) at a value of the metric. For example, building a CpG-context-based model allows us to compute the quantity P(Y=1 | CpG content), and then we consider the difference:

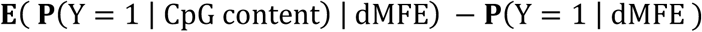

Squaring this difference and taking a weighted sum over all values of dMFE in a context, we recover the variance left unexplained by a particular non-structural variable. We obtain an R^2^ by comparing unexplained variance to that obtained using a null model, and the Depleter is then the variable with the largest R^2^ (see **Methods** for more details). **Table 1** shows that Depleters can recover large portions of the trends in **F****igure** **5** and **Supplementary FIGURES 4-5.** The striking trend between dMFE and gnomAD frequency in CpG-transitional contexts is largely driven by the proportion of CpGs in the surrounding 120 nucleotides (73% for CpG>TpG sSNVs and 79% for CpG>CpA sSNVs). CpG content is also the most powerful feature when accounting for the behavior of CFEED and dCD in these contexts, with high CpG content consistently correlating with depletion. The natural inference is that an abundance of CpGs signifies important mRNA structure nearby, the disruption of which could be deleterious.

In non-CpG-transitional contexts, the Depleter almost always proves to be a nucleotide upstream or downstream of the sSNV. In the context C>A we can recover 28% of the relationship between dMFE and gnomAD frequency simply by looking at whether the C is followed by a G. The power of CpG dinucleotides in recovering our structural trends in the contexts C>A, C>G, G>C and then G>T, emphasizes the powerful but poorly understood role of CpGs in both mRNA stability and mammalian genomes.

### Global quantification of mRNA constraint

Our analysis shows that variants predicted to influence mRNA secondary structures are constrained in the population. However, due to the multiple facets that need to be considered when studying RNA secondary structure, by focusing on a single RNA-folding metric such as dMFE or CFEED, we run the risk of missing functionally relevant information. To overcome this potential limitation of our RNA folding metrics, we set out to devise a more diversified method for predicting possible pathogenicity due to mRNA structure. Our strategy is to consider the additional statistical power bestowed by mRNA structure. In each of our 14 sequence contexts from **Table 1** we build two general logistic models for predicting MAF >0: a null model that uses the natural variables of sequence context, local nucleotide composition, transcript position and tRNA propensity, but NOT mRNA structure (***n***); and a structural model which also includes the 10 metrics obtained from our ViennaRNA analysis (***s***). These models yield two separate probability-predictions *P*_*n*_ and *P*_*s*_ for the quantity P(MAF >0) (see **Methods** for details). Then we define the metric:

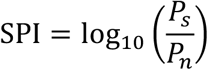

The metric SPI thus measures the additional predictive power bestowed by mRNA-structural variables. When it varies from 0, mRNA structural predictions yield new insight about a SNV’s potential to have a functional role in mRNA secondary structure. The power of SPI in each context (given by its area under the curve in predicting whether gnomAD is >0) is supplied in **Table 2** and we plot SPI vs. Y in CpG-transitional contexts in **F****igure** **6** (and in all contexts in **Supplementary FIGURE 6**). The classification rules of SPI vary widely by context. We see the most impressive performance in the context of CpG transitions. For both CpG>CpA and CpG>TpG transitions, those sSNVs with low SPI values are clearly under constraint.

**FIGURE 6.**
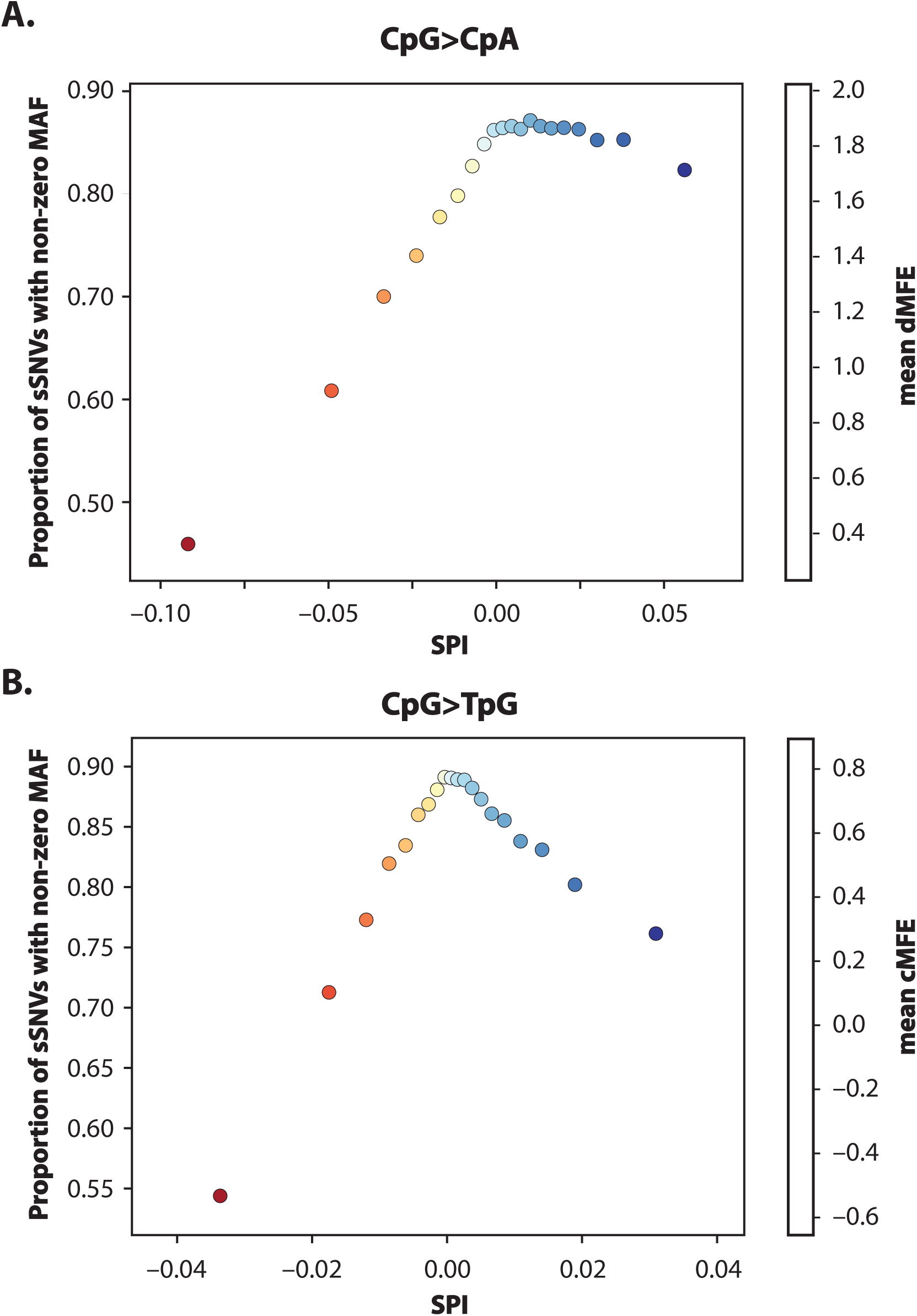
SPI score correlates with constraint in synonymous CpG transitions. Variants in the contexts **(A)** CpG>CpA and **(B)** CpG>TpG are divided by SPI score into 20 equal bins and the value P(MAF>0) plotted against the mean of each bin. We also colored by the mean dMFE over each bin. In both contexts the constraint is highest towards negative SPI, i.e. sSNVs for which structural information decreases the predicted probability that MAF > 0.

The behavior of SPI in non-CpG-transitional contexts is less regular and harder to weave into a coherent story. Every context shows a clear pattern, but this may amount to either enrichment or depletion (or both) as SPI moves in either direction. Given the strong dependence on REF-ALT context, the use of SPI as a deleteriousness score in non-CpG contexts may need further evaluation.

### Clinical Examples of Structural Pathogenicity

The literature reveals only a few examples of synonymous sSNVs unequivocally shown to be pathogenic through their effects on mRNA structure. These sSNVs, with accompanying values of our three ViennaRNA metrics and SPI, are listed in **Table 3.** The sSNVs show a definite enrichment for our structural metrics as each shows a value of |dMFE|, CFEED, |dCD| or |SPI| that is in at least the 80^th^ percentile in its context. For example, the pathogenic sSNV in NKX2-5, linked to congenital heart disease, has a dCD score in the 90^th^ percentile [44]. It should be noted that none of these clinical sSNVs qualify as truly exceptional outliers for any of our ViennaRNA metrics or SPI; all have scores below the 95^th^ percentile for |dMFE|, CFEED, |dCD| or |SPI| (see **DISCUSSION** for suggested score cutoff values).

## Discussion

We have shown that *in silico* mRNA structural predictions can be used to predict and explain the population allele frequency of a synonymous variant. By calculating ViennaRNA folding metrics for nearly 0.5 billion possible SNVs, we demonstrate that there is significant selection against sSNVs predicted to either stabilize or de-stabilize the given transcript’s local mRNA secondary structure. While the observed trends can be partially explained by sequence-based variables like CpG or GC content or membership in a CpG/AT/TA dinucleotide (as given by the “Depleter” field in **Table 1**), we believe our data support our hypothesis that RNA structure itself plays a critical role in human health and disease. As such, polymorphisms impacting mRNA structure are under negative selection in the population and should be more carefully evaluated in the context of both Mendelian disorders and complex human disease.

When determining if the connection between mRNA disruption and population incidence is direct and causal, we need to consider three factors. First, constraint of mRNA structure *must* be exercised through sequence-based variables, since the underlying primary mRNA sequence largely determines the secondary structure. Thus, although the trends we have observed may be influenced by sequence features such as CpG content (as illustrated by the “Depleter” variable in **Table 1**), it does not necessarily indicate that the trends are spurious. Second, it is important to note that our trends operate in the directions implied by our hypothesis: sSNVs that disrupt mRNA (measured three different ways) are depleted rather than enriched in the population for almost all REF>ALT contexts. Third, mutations that are predicted to change stronger base pairs to weaker ones are consistently constrained against de-stabilization rather than over-stabilization. If the association were spurious, we would not expect such agreement with prediction.

Our data also include some patterns which can be elegantly explained through mRNA structure. For example, when considering CFEED in **F****igure** **4****B** we observed that sSNVs were enriched when their CFEED values were multiples of 4. Since a CFEED value being divisible by 4 is a necessary condition to preserve the total number of base-pairs, this observed enrichment suggests changes to base pairing are constrained. We also observed bi-directional constraint for dMFE in the context T>C, visible in **Supplementary FIGURE 5** and also inferable from the low quadratic p-value in **Table 1**. We conjecture the dual constraint in this context might be due to guanine’s unique ability to wobble base-pair. Wobble base-pairing occurs between two nucleotides such as guanine-uracil (G-U), that are not canonical Watson-Crick base pairs, but have comparable thermodynamic stabilities. Thus, the dual constraint from mutations T>C could be related to the transformation of T=G wobble base-pairs into stronger C=G Watson-Crick base pairs.

Finally, in addition to the metrics output by our ViennaRNA analysis, we devised our own metric to measure structural pathogenicity. The Structural Predictivity Index (**SPI**), created specifically to control for all confounding factors, shows that mRNA structure has predictive power all by itself. Also, the clusters of outliers at the extreme values of our structural metrics (see **F****IGURES** **5** and **Supplementary FIGURES 4**,**5)** suggest a constraint beyond that explained by any confounding variable.

Taken together, this evidence provides significant support for the hypothesis that disruptions to mRNA structure are directly under constraint. However, we also realize there are other molecular mechanisms at work in the regulation of the transcriptional and translational processes. For example, while the retention of CpG dinucleotides is certainly connected to mRNA structure, other factors such as regulatory methylation, tRNA binding, binding of miRNAs and other RNA binding proteins, DNA chromatin structure and epigenetic modifications in the ORF could also be involved. Relatedly, we found a few contexts where sSNVs which disrupt mRNA structure actually have higher population frequencies than those that do not (**Table 1)**, the main example being the enrichment of high CFEED values in the contexts C>A and G>A. In both cases, the Depleter variable is a trailing A, which correlates negatively with gnomAD frequency and CFEED. The presence of an A is likely to have minimal effect on mRNA structure, suggesting that in these contexts the connection is partly spurious.

### Successful identification of structurally disruptive sSNVs in known pathogenic synonymous variants

Over the last decade numerous studies have demonstrated that synonymous variants play essential molecular roles in regulating both mRNA structure and processing, including regulation of protein expression, folding and function [reviewed in 12, 60, 61]. However, the potential for pathogenic synonymous variants that impact RNA folding in human genetic disease remains largely unknown. Current American College of Medical Genetics (**ACMG**) guidelines for the assessment of clinically relevant genetic variants focus primarily on missense, nonsense or canonical splice variants [9]. These guidelines suggest that synonymous “silent” variants should be classified as likely benign, if the nucleotide position is not conserved and splicing assessment algorithms predict neither an impact to a splice consensus sequence nor the creation of a new alternate splice consensus sequence. In the absence of functional tools that would aid in the simultaneous assessment of both nsSNVs and sSNVs in a given patient’s genome, we are almost certainly missing novel disease etiologies that have their molecular underpinnings in pathological alterations to mRNA structure.

Numerous *in silico* tools exist to aid in the prediction of disease-causing missense variants, and have accuracy in the 65%-85% range when evaluating known pathogenic variants [62]. Such algorithms infer pathogenicity based on amino-acid substitutions (SIFT [63], PolyPhen [64], FATHMM [65]), nucleotide conservation (SiPhy [66], GERP++ [67]) or an ensemble of annotations and scores (CADD [68], DANN [69], REVEL [70]). These tools predict whether a nsSNV is pathogenic or benign, primarily due to the high conservation of protein sequences. However, these algorithms are not equipped to assess pathogenicity in synonymous variants, which are under different constraints [11]. Recognizing that there is a critical need for methods that better predict the potential whether sSNVs have pathogenic impact and function, our goal in this present study was the generation of such metrics. ViennaRNA stability and SPI metrics are available for download for all known sSNVs, to enable researchers and clinicians to evaluate WES and WGS data in combination with tools such as Annovar [71], SnpEff [72] and VEP [73].

The role of synonymous mutations in human disease is gaining recognition in the community. The RNAsnp Web Server predicts the change in optimal mRNA structure and base-pairing probabilities due to a SNV [74]. A command line tool called remuRNA calculates the relative entropy between the mutant and wildtype mRNA structural ensembles [75]. While these tools predict disruptions to mRNA structure, they do not attempt to predict pathogenicity and must be executed manually on each variant of interest. Both RNAsnp and remuRNA were recently utilized to create a database of synonymous mutations in cancer (SynMICdb), using data from COSMIC across 88 tumor types [76]. For constitutional genetic disease, a related resource is the databases of Deleterious Synonymous (dbDSM), which manually curates sSNVs reported to pathogenic in the literature and in databases like ClinVar [77]. These resources represent an important step towards evaluating sSNVs in disease. However, relatively few sSNVs have well supported evidence of their pathogenicity, due in part to the complexity of demonstrating the molecular impact of these variants through functional studies. While in the above databases and elsewhere we found dozens of examples of sSNVs suggested to be pathogenic due to their effects on mRNA structure, most of these cases lacked functional studies to conclusively implicate the given variant in disease. As such we focused on a set of six sSNVs that we believe the authors unequivocally demonstrated to be pathogenic through their effects on mRNA structure (**Table 3**). This dataset included one variant in OPTC associated with glaucoma [42], two variants in NKX2-5 associated with congenital heart defects [44], one variant in DRD2 associated with post-traumatic stress disorder [40], and two variants in COMT associated with pain sensitivity [41].

All six sSNVs demonstrated definite enrichment for our structural metrics, by stability, edge distance, diversity or SPI, with values in the 80^th^ to 90^th^ percentile range. However, none of these clinically relevant sSNVs qualifies as a truly exceptional outlier for any of our ViennaRNA metrics or SPI with all percentiles being below 90. It is theoretically possible that such extreme outliers are not biologically tenable, making them less likely to appear in the human population. As such, a change in the 80^th^ percentile could represent a cutoff for biological significance. Another possibility (perhaps equally strong) is that these sSNVs occupy important regulatory positions, and that a sSNV deleterious to mRNA secondary structure may exhibit pathogenicity when it distorts structure *in a key region* of the transcript.

The enrichment of our structural metrics, while moderate, is still clear and our hope is that future studies will allow refinement and enhancement of our metrics. As new discoveries of pathogenic sSNVs in human genetic disease occur, a larger data set of known clinically relevant sSNVs will help determine cutoff values. For now, our recommendation is that a conservative 80^th^ percentile cutoff across the four metrics is used initially, but this may need to be lowered to reveal pathogenic sSNVs that have a less extreme change to mRNA structure.

### Molecular mechanisms underlying constraint of sSNVs

Synonymous variants that impact mRNA secondary structure could confer pathogenicity in numerous ways. For example, sSNVs can alter global RNA stability, where less stable RNA molecules may be degraded more quickly resulting in lower protein levels [23, 25, 27]. As local RNA structure is essential for the translation process, a more stable mRNA may not be able to initiate translation, also resulting in lower protein levels [20, 21, 30, 31]. Additionally, numerous studies argue that by making the mRNA structure too difficult, or too easy, for the ribosome to process, synonymous codons may act as a subliminal code for protein folding [16, 30, 32, 33, 35].

While there is a large body of literature supporting the essential role of sSNVs in preservation of mRNA secondary, sSNVs also play roles in other molecular processes that could impact our observations. While the stability of an mRNA transcript can determine how quickly it is translated [19, 29, 30], protein synthesis is also regulated by both the abundance [78] and recruitment of tRNAs through synonymous codon utilization (codon bias) [79-81]. Changes in the bicodon bias has the potential to lead to pathogenicity of synonymous mutations in human disease [16, 35]. As such, this may have had a confounding impact on our analysis of constraint, but we attempted to mitigate this by including the tRNA Adaptivity Index (a measure of tRNA abundance) in our set of confounding variables.

Some of our constraint might have its roots in the tertiary DNA structure, where DNA bending plays roles in DNA packaging, transcriptional regulation, and specificity in DNA–protein recognition [82]. While it has been suggested that synonymous codon choice may play a role in DNA bending [83], the process is largely governed by intergenic and intronic DNA, and as such is unlikely to be directly responsible for the constraint shown in **F****igure** **2**. We attempted to mitigate the role of local DNA sequence context and CpG content, by including both in our set of confounding variables when generating SPI. However, methylation of CpG dinucleotides can regulate gene expression, potentially making it difficult to state whether selection for mRNA structure causes the retention of CpGs, or whether the retention of CpGs is regulated by a process independent of mRNA structure. A strong reason for CpGs to operate causally with regard to mRNA structure is that they are the most important determinant of mRNA structure (along with G+C content, with which CpG content correlates closely). Retention of 5’ ORF CpG sites occurs at a high frequency in the first exon of coding genes; a stacked C:G + G:C base pairing has the lowest free energy of the 36 possible stacked base-pair combinations [84]; and deamination of CpGs can be suppressed and repaired by existing enzymatic mechanisms. Thus, CpG dinucleotides represent the easiest and most natural way to determine mRNA structure.

Finally, it is important to also consider the essential role that synonymous variants play in RNA splicing. While we took care to exclude sSNVs impacting the canonical splice sites from our constraint analysis, exonic variants beyond the canonical splice site can disrupt splice enhancers [85], or they may also activate cryptic splice sites, leading to aberrant pre-mRNA splicing and loss of coding sequence [86]. Given the diversity of molecular roles that synonymous codons have, it will be important for future studies to create scores that would allow assessment of sSNV pathogenicity through any these possible mechanisms.

## Conclusions

We have shown that sSNVs which stabilize or destabilize mRNA are significantly constrained in the human population, thereby supporting a growing body of evidence that previously assumed “silent” polymorphisms actually play crucial roles in regulation of gene expression and protein function. We have demonstrated that this connection is rich, complex, and biologically intuitive. Given that there are multiple mechanisms by which sSNVs influence biological function, we are almost certainly missing undiscovered disease etiologies when these variants are ignored. In addition to providing the community with a dataset of ten ViennaRNA structural metrics for every known synonymous variant, our Structural Predictivity Index is the first metric of its kind to enable global assessment of sSNVs in human genetic studies. We hope that these metrics will be utilized to accurately assess and prioritize an underrepresented class of genetic variation that may be playing significant and as yet to be realized role in human health and disease.

## Methods

### Raw Dataset

To obtain all human mRNA transcripts we downloaded the NCBI RefSeq Release 81 from an online repository (ftp://ftp.ncbi.nlm.nih.gov/refseq/H_sapiens/mRNA_Prot/). Transcript sequences corresponded to human reference genome build GRCh38.

### Overview of RNA structure prediction process

To estimate the structural properties of a sSNVs we used the ViennaRNA software package, a secondary structure prediction package that has been extensively utilized and continuously developed for nearly twenty-five years. ViennaRNA uses the standard partition-function paradigm of RNA structural prediction [87]. We utilize version 2.0 of ViennaRNA [18]. Applying ViennaRNA to every possible SNV in the human genome (about 500,000,000 calculations) was a computationally challenging task which we carried out using an Apache Spark framework powered by Amazon Web Services (AWS). We built a pipeline which read in and analyzed a SNV and stored the results in AWS Simple Storage Service (S3) in Parquet columnar file format (**F****igure** **2**). The ease and capacity of AWS greatly facilitated the project, and the affordability of S3 storage means our data can easily be shared with others. The software we developed is available on GitHub: https://github.com/nch-igm/rna-stability.

### RNA structure prediction methodology

To analyze a given SNV we built a 101-base sequence consisting of a central nucleotide at the 51st position (which we set to either the reference or the three alternates) along with the 50 flanking bases on either side. If the nucleotide lay 50 bases from the transcript boundary, the window was simply taken to be the first or last 101 bases in the transcript. We processed these sequences in fasta format with ViennaRNA’s flagship module RNAfold, which yielded three predicted mRNA secondary structures – the minimum free energy, centroid, and maximum expected accuracy structure – as well as numeric values for the free energy of each structure, and a fourth metric measuring the energy of the whole ensemble (see the documentation of [18] for detailed descriptions of these concepts). Comparing the free energies between the wildtype and mutant for each type of structure gave us the four stability metrics delta-MFE (dMFE), dCFE, dMEAFE and dEFE. Next, the predicted structures were processed by the ViennaRNA module RNApdist, which counted the edge-differences to produce the four edge-metrics MFEED (minimum free energy edit distance), CFEED, MEAED and EFEED. As a final step, the predicted structures were further processed by the ViennaRNA program RNAdistance to obtain the diversity metrics dCD and dEND (change in distance from centroid and change in ensemble diversity, respectively).

This whole procedure was carried out using custom developed Spark wrappers of RNAfold, RNApdist and RNAdistance, with slight modifications to the source code to suppress the creation of graphics files. After building our fasta files, we were able to compute all 10 ViennaRNA metrics for over a half billion sequences in less than 24 hours using 51 c4.8xlarge AWS EMR computing nodes.

### Construction of final dataset for synonymous SNVs

The next step was to extract the sSNVs. This task was complicated by the fact that a SNV might have appeared in several different transcripts, and could be synonymous in some and non-synonymous in others. To address this challenge, we first annotated every SNV using the program snpEff [72], whose source code was modified to allow record-by-record calling via Spark. This snpEff analysis produced annotations of predicted biotype, e.g. missense, synonymous, canonical splice site, etc. To validate these snpEff predictions we manually predicted the biotype of each SNV using start and stop codon information from RefSeq (ftp://ftp.ncbi.nih.gov/refseq/H_sapiens/RefSeqGene/refseqgene.*.genomic.gbff.gz). The small number of sSNVs where our predicted biotype disagreed with snpEff’s were discarded. We then defined a “synonymous SNV” to be one that was (A) synonymous in at least one transcript, (B) synonymous or within the UTRs in all transcripts, and (C) not implicated in splicing by snpEff. Each sSNV identified as “synonymous” by this scheme was assigned a “home transcript,” chosen based on proximity to the start codon, then on maximal transcript coding sequence length, and then arbitrarily.

This filtration and duplicate-removal process yielded a final set of 17.9 million sSNVs in 34,000 transcripts. See **Supplementary Table 2** for a table giving the landscape of our final dataset and the number of sSNVs filtered at each stage.

### Merging of sSNV GRCh38 transcript coordinates with gnomAD GRCh37 coordinates

To measure constraint operating on a sSNV we used population frequencies obtained from the gnomAD database. Since this resource only existed for the GRCh37 reference build, we lifted our entire dataset from GRCH38 to GRCh37. The lifting procedure was carried out using the Picard Tools program liftOver [88], which was executed using a custom Spark wrapper. The joining of the gnomAD frequencies to our main dataset was a task greatly facilitated by Spark’s parallel processing and native Parquet support. Since the great majority (approximately 90%) of sSNVs were marked with gnomAD frequency 0, it is important to identify sSNVs marked zero purely through a lack of coverage. To achieve this, we flagged and removed all sSNVs where fewer than 70% of samples had at least 20X coverage.

### Further variant annotations

Next, we estimated the local nucleotide content around each sSNV. We divided each transcript into windows of 40 bases and in each window computed the proportion of A’s, C’s, G’s, T’s, CpG’s and AT’s in the surrounding three windows. Finally we joined multiple additional annotations (including conservation metrics such as PhyloP) from the dbNSFP dataset [89]. Again, this heavy task was greatly facilitated by our Spark framework.

### Partition of dataset

We carried out most of the analysis separately on subsets of data defined by a common mRNA reference and alternate allele, e.g those sSNVs of form C>A. The reference and alternate alleles exert such a huge influence on gnomAD frequency that the best solution seemed to be to control for them explicitly. Dividing our dataset based on mRNA alleles (as opposed to DNA alleles, which do not depend on transcript sense) is a step justified in **Supplementary Table 3**.

### Depleter variables

Depleter variables (so called because they explain some of the gnomAD depletion at values of a structural variable) are given in **Table 1**. They are chosen to be the sequence feature that explains the greatest portion of the connection between a structural metric (e.g. dMFE) and Y in a context. Possible Depleter variables are local nucleotide content and the specific nucleotides up/downstream of the sSNV.

To compute the correlation between a structural metric (e.g. dMFE) and Y that is left unexplained by a sequence feature (e.g. CpG content) in a particular REF-ALT context, we first build a simple logistic regression model between CpG content and Y, which gives us an estimate **P**(Y = 1 | CpG content) for every sSNV in the context (based on the proportion of CpGs in the surrounding 120 nucleotides). We then plug this “structure-less” estimate into the expression

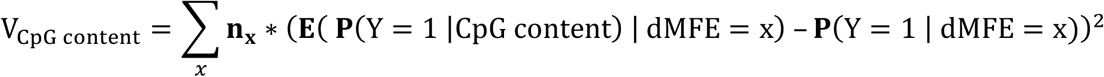

where we sum over all values x of dMFE and let **n**_**x**_ denote the number of sSNVs in the context with dMFE= x. Comparing this quantity to the null variance

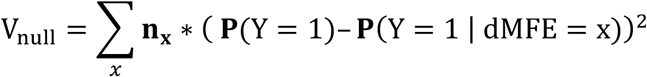

allows us to compute the proportion of the variation explained by CpG content:

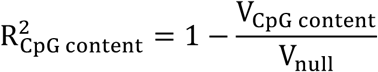

The “Depleter” for a given structural metric in a given context is chosen as the variable with the highest R^2^. Finally the correlation between the Depleter and Y was checked, and the Depleter given a sign (+/-) so that the signed Depleter correlated negatively with Y.

### Construction of SPI

To construct our final SPI scores we built two separate models over each of our 14 contexts to predict the event MAF > 0. The “null” model used all natural features -the nine nucleotides in the SNV’s home and adjacent codons, the proportion of A/C/G/T/CpG/AT’s in the surrounding 120 nucleotides, the sSNV’s position in its transcript and the transcript’s length, and the tAI (tRNA Adapation Index obtained from a supplement of [90] from https://ars.els-cdn.com/content/image/1-s2.0-S0092867410003193-mmc2.xls) of the wildtype and mutant codons. The second, “active” model used all these features plus our 10 ViennaRNA metrics. Both sets of variables were then used to predict MAF > 0. We then defined the SPI score for a sSNV to be the base-10 logarithm of the active model’s predicted P(Y=1) probability divided by the null model’s predicted P(Y=1). Context wise plots for SPI are given in the **Supplementary Figure 6**.

We tried three different model-styles for computing the raw predictions that comprise SPI – general logistic as implemented in python’s sklearn LogisticRegression module, random forest as implemented in sklearn’s RandomForestClassifier, and gradient-boosted trees as implemented in the python package xgboost. Performance of each SPI “flavor” is given in **Supplementary Table 4**. We eventually settled on the general logistic model, as it out-performs the gradient-boosted tree model and does not over-fit as the random forest mode does.

## Declarations

### Ethics approval and consent to participate

Not applicable.

### Consent for publication

Not applicable.

### Availability of data and materials

The software we developed and structural scores are available on GitHub: https://github.com/nch-igm/rna-stability.

### Authors’ contributions

J.B.S.G., J.L.L and P.W. developed methodology, performed data analysis and results interpretation. G.E.L. developed AWS Spark ViennaRNA pipeline and developed variant annotation tools. G.E.L. generated folding metrics. J.B.S.G. developed Structural Predictivity Index (SPI). D.M.G., H.C.K., B.J.K, and J.R.F assisted with data analysis, interpretation of results and development of variant annotation tools. J.B.S.G, G.E.L and P.W. prepared figures. All authors contributed to the preparation and editing of the final manuscript.

## Acknowledgements

This team works in the Steve and Cindy Rasmussen Institute for Genomic Medicine at Nationwide Children’s Hospital. The Institute is generously supported by the Nationwide Foundation Pediatric Innovation Fund.

## Additional Files

**S****upplementary** **D****ata** **F****ILE** **1**: This file contains four supplementary data tables and six supplementary figures further detailing the methodology and results presented in this manuscript.

### FIGURE LEGENDS

**Table 1. Structural metrics correlate with gnomAD frequency in most REF>ALT contexts.** Correlation between structural metrics (**A**) dMFE, (**B**) CFEED and integer-rounded (**C**) dCD on the one hand, and the quantity P(MAF>0) on the other, over all sSNVs in a given context. The R^2^ and p-values are obtained from a weighted least-squares linear regression, with the p-value corresponding to the linear coefficient; a quadratic regression was also performed, but only the p-value was retained. Only context-metric pairs with p-value < 0.005 are included. “Normalized slope” was obtained by dividing slope of regression line by average P(MAF>0) in the context and then multiplying by range covered by metric in its central 90% of sSNVs. “Depleter” is raw sequence variable that explains largest proportion of structural trend *in this context*, with sign adjusted to correlate negatively with gnomAD frequency. “Depleter R^2^” gives proportion of variance explained by Depleter (see *Depleter variables* in **R****esults** for details).

**Table 2. Area under curve for SPI score.** SPI was used to discriminate MAF > 0 using a simple logistic model with 5-fold cross-validation. Table shows area under curve for model, averaged over the 5 training and testing sets.

**Table 3: Known sSNVs clinically implicated for structural pathogenicity are successfully predicted to be pathogenic by our structural metrics.** dbSNP RS number and standardized SNP annotations are provided, along with the genes official gene symbol and disease the sSNV has been associated with. The absolute value of dMFE, CFEED, dCD and SPI are provided, along with the percentile value of that score, computed over each context, in parentheses.

