## Supplemental Data for "Global Analysis of Human mRNA Folding Disruptions in Synonymous Variants Demonstrates Significant Population Constraint"

* Corresponding author

Mailing address:

The Institute for Genomic Medicine

Nationwide Children's Hospital

575 Children's Crossroad

Columbus, OH 43215. USA

**Keywords:** synonymous mutation, mRNA stability, RNA folding, RNA secondary structure, genetic disease, Spark, gnomAD, transcriptomics

SUPPLEMENTARY DATA TABLE 1

| Vienna metric | Vienna metric abbreviation | Description |
|---|---|---|
| ***Stability Metrics*** | | |
| **dMFE** | delta Minimum Free Energy | Change in stability of optimal mRNA structure |
| **dCFE** | delta Centroid Free Energy | Change in stability of centroid mRNA structure |
| **dMEAFE** | delta Maximum Expected Accuracy Free Energy | Change in stability of maximum expected accuracy mRNA structure |
| **dEFE** | delta Ensemble Free Energy | Expected change in stability over all structures |
| ***Edge distance metrics*** | | |
| **MFEED** | Minimum Free Energy Edit Distance | Edge-changes in optimal structures |
| **CFEED** | Centroid Free Energy Edit Distance | Edge-changes in centroid structures |
| **MEAED** | Maximum Expected Accuracy Edit Distance | Edge-changes in maximum expected accuracy structures |
| **EFEED** | Ensemble Free Energy Edit Distance | Expected edge-changes over all structures |
| ***Diversity metrics*** | | |
| **dCD** | delta Centroid Distance | Change in average distance of structural ensemble from centroid structure |
| **dEND** | delta Ensemble Diversity | Change in ensemble diversity |

**Supplementary Data Table 1. Description of the ten Vienna RNA metrics calculated through the Spark RNA stability pipeline.** These metrics are divided into three classes: stability, edge distance and diversity.

SUPPLEMENTARY DATA TABLE 2

| Variant type | Total # SNVs, counted by transcript position | Total # SNVs, counted by chromosomal position | # SNVs successfully lifted to hg19 | # SNVs passing gnomAD filter | # SNVs not implicated in splicing | # SNVs with adequate gnomAD coverage | Final # SNVs |
|---|---|---|---|---|---|---|---|
| **all** | 469,071,297 | 203,647,788 | 203,430,738 | 201,889,552 | 200,292,238 | 185,372,924 | 181,809,836 |
| **5′ utr** | 33,684,299 | 15,242,049 | 15,226,793 | 15,191,485 | 15,038,625 | 15,242,049 | 14,988,664 |
| **3′ utr** | 208,223,247 | 85,572,534 | 85,522,731 | 85,365,605 | 85,387,182 | 85,572,534 | 85,180,818 |
| **missense** | 161,898,807 | 72,878,682 | 72,772,402 | 71,904,594 | 70,700,280 | 59,831,999 | 57,738,594 |
| **stopgain** | 9,288,276 | 4,155,868 | 4,150,780 | 4,113,672 | 4,016,853 | 3,465,335 | 3,338,052 |
| **stoploss** | 349,722 | 177,085 | 176,929 | 175,711 | 170,495 | 122,062 | 117,943 |
| **startgain** | 4,491,945 | 2,475,690 | 2,473,076 | 2,467,518 | 2,475,690 | 2,475,690 | 2,467,518 |
| **startloss** | 407,050 | 226,666 | 226,450 | 222,459 | 218,757 | 127,608 | 121,970 |
| **synonymous** | 50,727,951 | 22,919,214 | 22,881,577 | 22,448,508 | 22,284,356 | 18,535,647 | **17,856,277** |

**Supplementary Data Table 2 summarizes data pre-processing steps.** Vienna metrics were calculated for a total of 469,071,297 SNVs in all known transcripts. As multiple transcripts share the same exonic genomic coordinates, we first collapsed the data to 203,647,788 unique chromosome positions, letting assigning each variant the most damaging variant-type represented at a position (the majority of duplicate positions represented only one variant-type). We then filtered out variants that could not be lifted from GRCh38 to matching hg19 (GRCh37) coordinates; that were flagged by gnomAD as having an unreliable population frequency; that lacked adequate gnomAD coverage; and that were predicted by SnpEff predicted to play a role in splicing. Finally, variants marked as “synonymous” (meaning they were synonymous in some transcript, and either synonymous or UTR in every transcripts) were extracted, giving us a core data set of 17,856,277 sSNVs.

SUPPLEMENTARY DATA TABLE 3

| mRNA REF allele | mRNA ALT allele | Transition (Ti) / Transversion (Tv) | # potential synonymous variants | P(MAF>0): + strand | P(MAF>0): - strand |
|---|---|---|---|---|---|
| A | G | Ti | 1,754,361 | 0.137 | 0.135 |
| A | C | Tv | 1,212,199 | 0.040 | 0.040 |
| A | T | Tv | 986,499 | 0.031 | 0.032 |
| C | T | Ti | 2,650,886 | 0.173 | 0.172 |
| C | G | Tv | 1,277,107 | 0.074 | 0.073 |
| C | A | Tv | 1,607,564 | 0.055 | 0.055 |
| G | A | Ti | 1,983,179 | 0.159 | 0.161 |
| G | T | Tv | 985,565 | 0.066 | 0.064 |
| G | C | Tv | 998,787 | 0.062 | 0.062 |
| T | C | Ti | 2,311,979 | 0.115 | 0.116 |
| T | G | Tv | 980,595 | 0.037 | 0.038 |
| T | A | Tv | 1,138,567 | 0.026 | 0.026 |
| **CpG Sequence Context** | | | | | |
| C | T | Ti | 283,968 | 0.825 | 0.831 |
| G | A | Ti | 172,742 | 0.801 | 0.806 |

**Supplementary Data Table 3 gives number of potential and actual synonymous variants in each context.** In each mRNA context we give the total number of potential synonymous variants in the human genome (subject to the filters imposed in Supplementary Data Table 2) as well as the proportion of these potential sSNVs that appear in gnomAD (i.e. MAF >0). We compute separate proportion over positive-sense and negative-sense home transcripts. The close agreement of the values of P(MAF>0) between positive- and negative-sense strands justifies our decision to classify by RNA rather than DNA context.

SUPPLEMENTARY DATA TABLE 4

| Context | GLM AUC Training Dataset | GLM AUC Testing Dataset | XGB AUC Training Dataset | XGB AUC Testing Dataset | RF AUC Training Dataset | RF AUC Testing Dataset |
|---|---|---|---|---|---|---|
| **A>C** | 0.521 | 0.516 | 0.499 | 0.491 | 0.775 | 0.513 |
| **A>G** | 0.539 | 0.539 | 0.515 | 0.512 | 0.847 | 0.545 |
| **A>T** | 0.512 | 0.505 | 0.495 | 0.484 | 0.714 | 0.514 |
| **C>A** | 0.550 | 0.548 | 0.505 | 0.501 | 0.758 | 0.536 |
| **C>G** | 0.513 | 0.509 | 0.519 | 0.526 | 0.776 | 0.540 |
| **C>T** | 0.506 | 0.506 | 0.509 | 0.510 | 0.870 | 0.511 |
| **CpG>TpG** | 0.656 | 0.656 | 0.509 | 0.510 | 0.902 | 0.614 |
| **G>A** | 0.522 | 0.524 | 0.514 | 0.513 | 0.869 | 0.511 |
| **CpG>CpA** | 0.616 | 0.618 | 0.513 | 0.514 | 0.896 | 0.621 |
| **G>C** | 0.536 | 0.538 | 0.524 | 0.526 | 0.772 | 0.540 |
| **G>T** | 0.534 | 0.532 | 0.531 | 0.536 | 0.701 | 0.548 |
| **T>A** | 0.540 | 0.532 | 0.495 | 0.484 | 0.581 | 0.502 |
| **T>C** | 0.565 | 0.564 | 0.507 | 0.507 | 0.844 | 0.544 |
| **T>G** | 0.529 | 0.522 | 0.498 | 0.483 | 0.787 | 0.507 |

**Supplementary Data Table 4 gives performance of SPI score under different model frameworks.** In each context we test the power of an SPI score built under one of three different schemes (general logistic, random forest, and gradient-boosted trees) in predicting whether a sSNV has MAF>0. Metric AUC measures the area under the receiver operating characteristic curve, averaged over a 5-fold cross validation. We ultimately use the general logistic mode (glm), since the gradient-boosted trees model (xgb) is not as powerful and the random-forest model (rf) is prone to overfitting (with the training AUCs being much greater than the testing AUCs).

### SUPPLEMENTARY DATA FIGURE 1 - Distribution of Structural Metrics

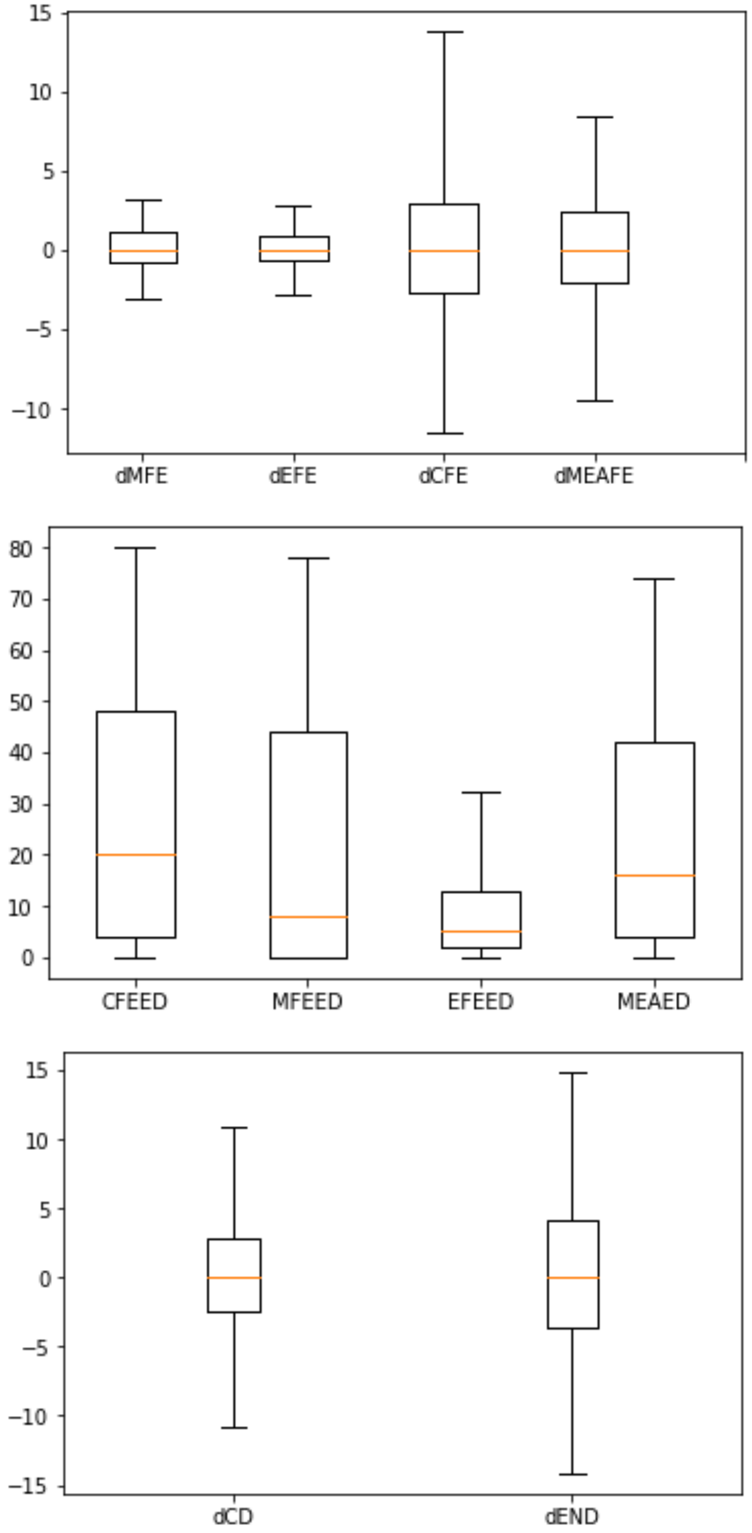

**Supplementary Data Figure 1. Global distribution of structural metrics using box-and-whisker plots.** We show the distribution of each metric our filtered database of possible sSNVs. Orange line shows median and box encloses central 75% of data. Whiskers enclose central 90%.

SUPPLEMENTARY DATA FIGURE 2 - Calculation of Edit Distance

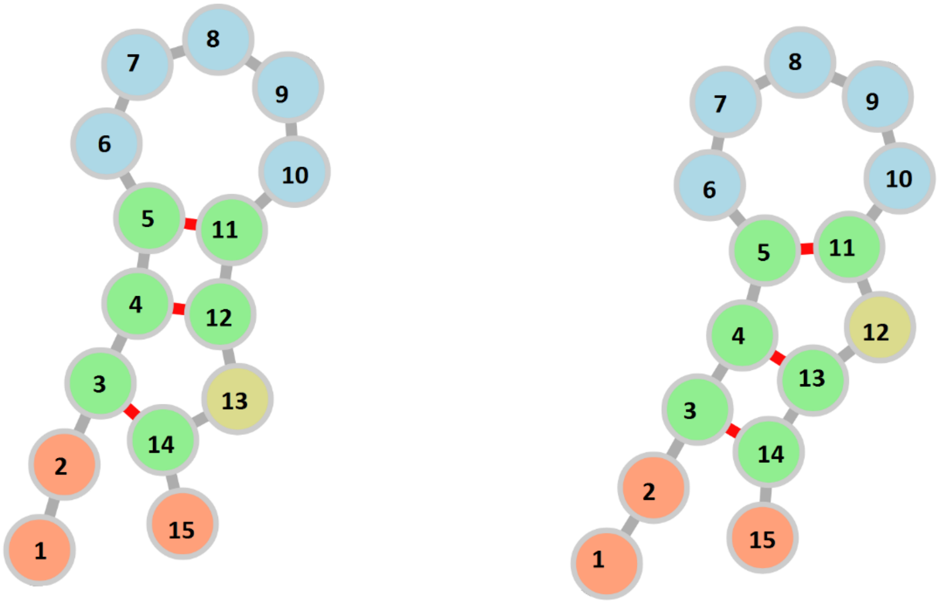

**Edit distance = 2 (#4 and #12 unpair) = 4**
**+ 2 (#4 and #13 pair)**

**Supplementary Data Figure 2. Calculation of edit distance.** The "edit distance" between two mRNA secondary structures with the same primary structure is the number of "edits" needed to transform one structure into another. Creation and removal of base pairs (the only possible changes) each count for two edits.

### SUPPLEMENTARY DATA FIGURE 3 - Global plots of structural metrics vs. Y

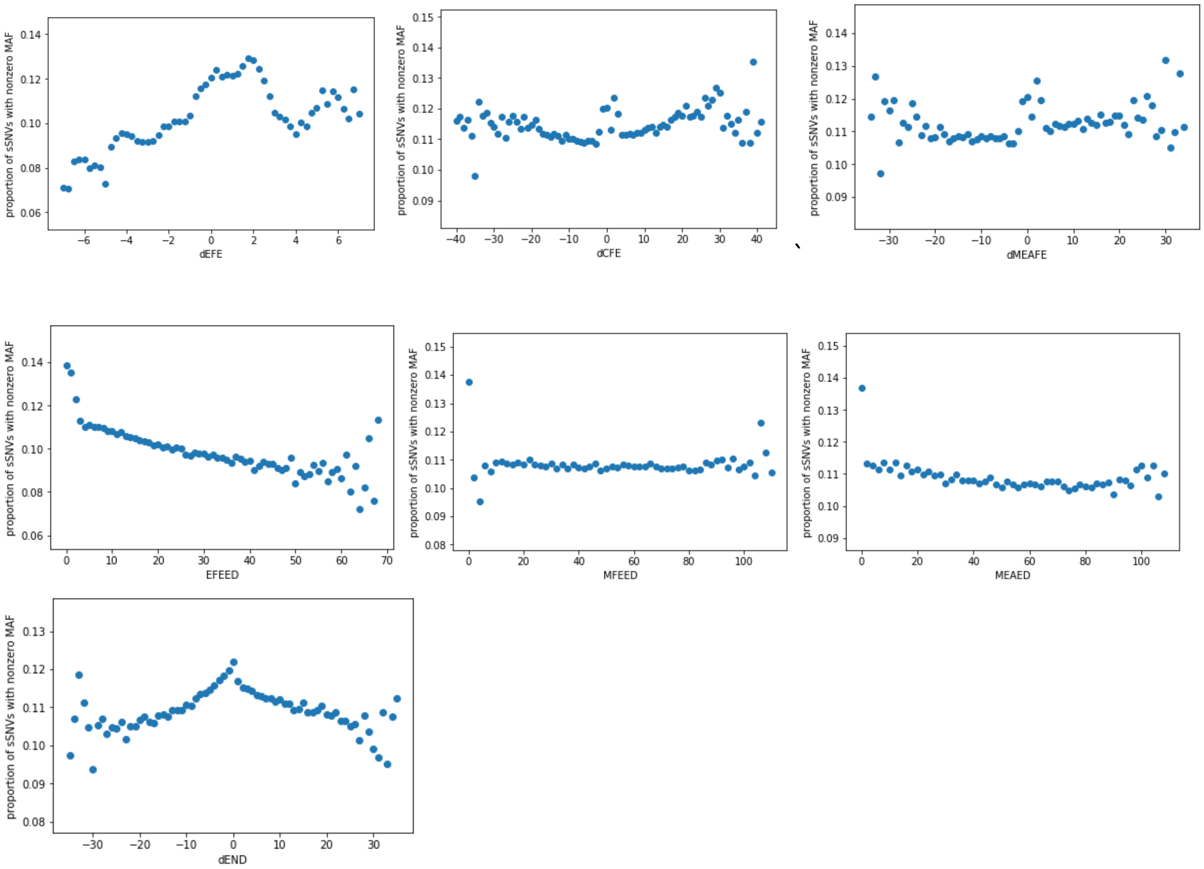

**Supplementary Data Figure 3. Vienna metrics vs. Y.** For seven Vienna metrics not featured in main results, we plot P(Y=1) vs. metric value. We rounded dCFE, dMEAFE, EFEED and dEND to the nearest integer, and dEFE to the nearest 0.25, prior to plotting. Values with fewer than 50 sSNVs in gnomAD have been removed. Descriptions of metrics are given in **Supplementary Data Table 1**.

### SUPPLEMENTARY DATA FIGURE 4 - Structural metric plots in contexts constrained against destabilization

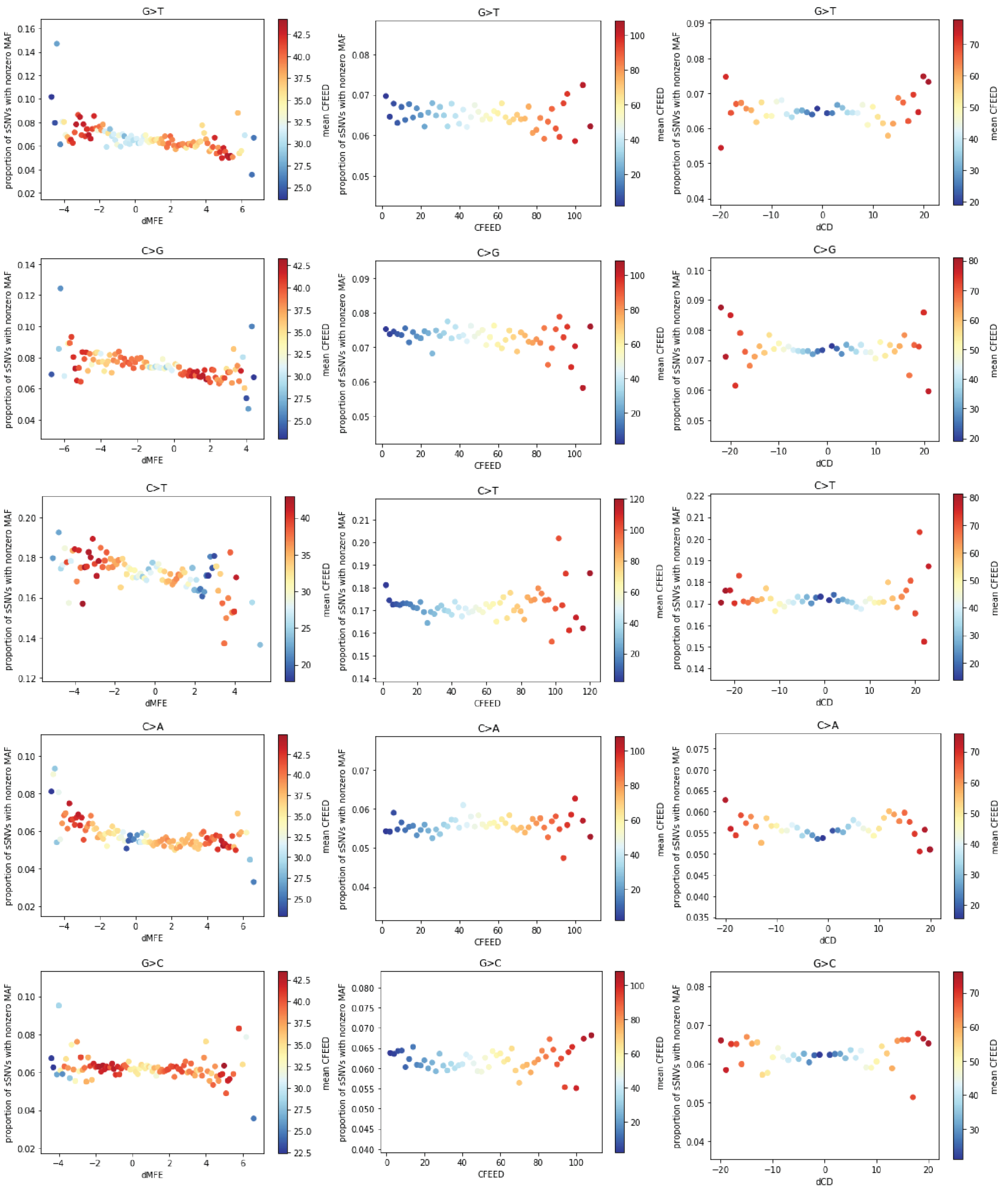

**Supplementary Figure 4. Primary Vienna metrics in contexts constrained against de-stabilization.** For every non-CpG-transitional context shown in Table 1A with a negative normalized slope (i.e. constraint against de-stabilization), we plot P(Y=1) vs. our three main Vienna metrics (dMFE, CFEED, dCD). Values of dCD were founded to the nearest integer prior to computing P(Y=1). Metric-values with fewer than 50 sSNVs in gnomAD are not shown.

**SUPPLEMENTARY DATA FIGURE 5 - Structural metric plots in contexts constrained against over-stabilization**

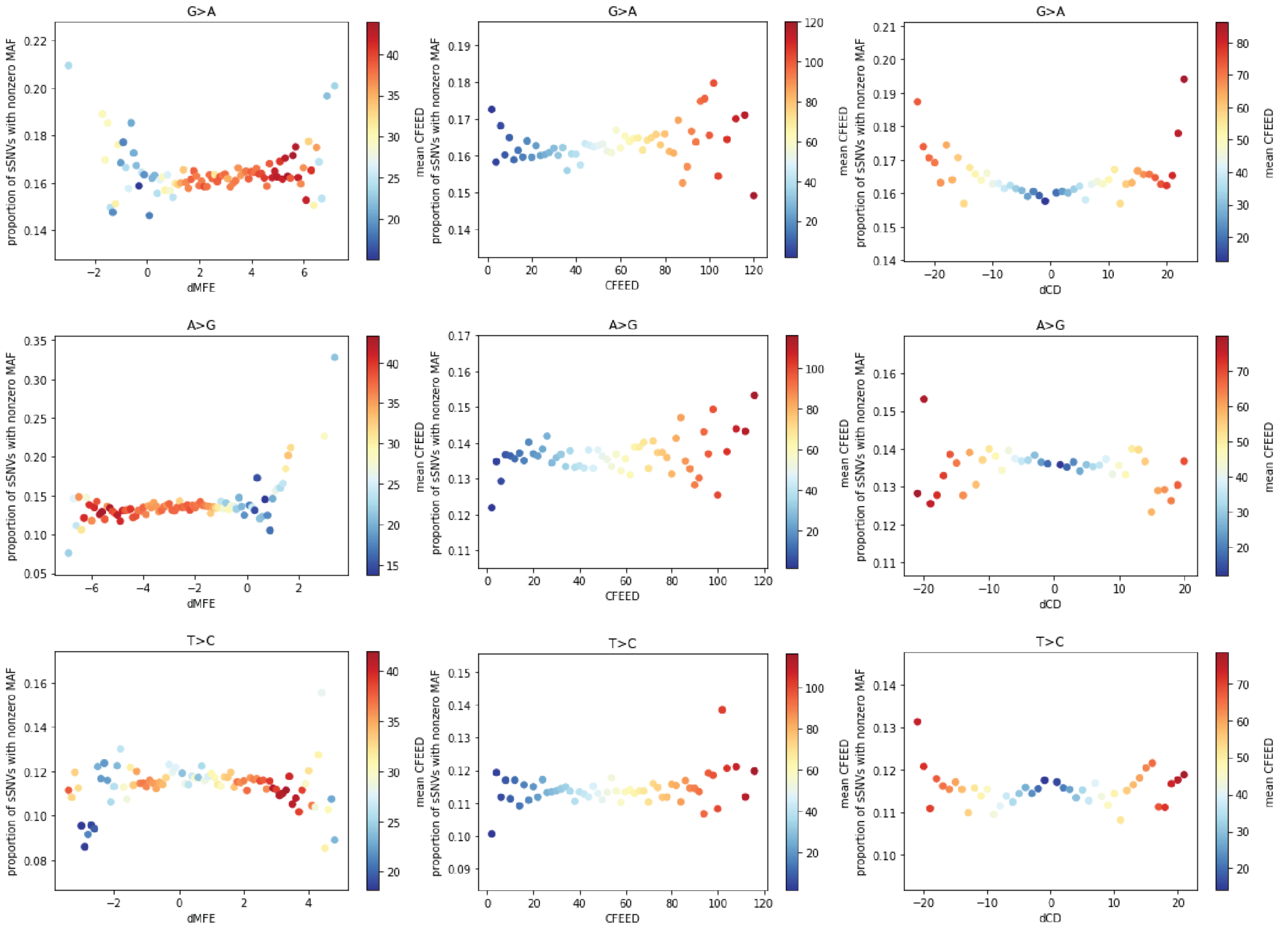

**Supplementary Data Figure 5. Primary Vienna metrics in contexts constrained against over-stabilization.** For every non-CpG-transitional context shown in Table 1A with a positive normalized slope (i.e. constraint against over-stabilization), we plot P(Y=1) vs. our three main Vienna metrics (dMFE, CFEED, dCD). Values of dCD were founded to the nearest integer prior to computing P(Y=1). Metric-values with fewer than 50 sSNVs in gnomAD are not shown.

### SUPPLEMENTARY DATA FIGURE 6 – Sequence Context and SPI

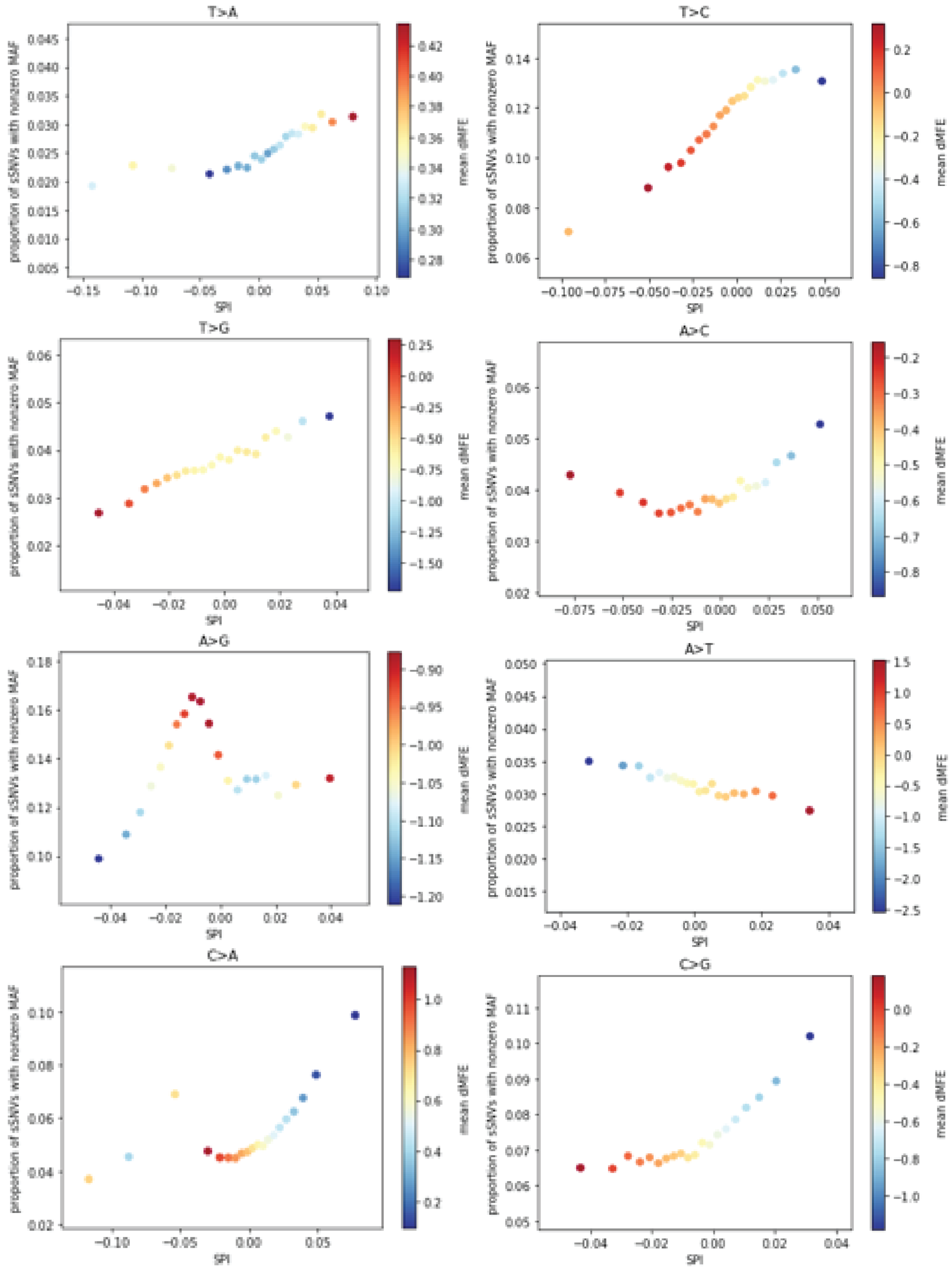

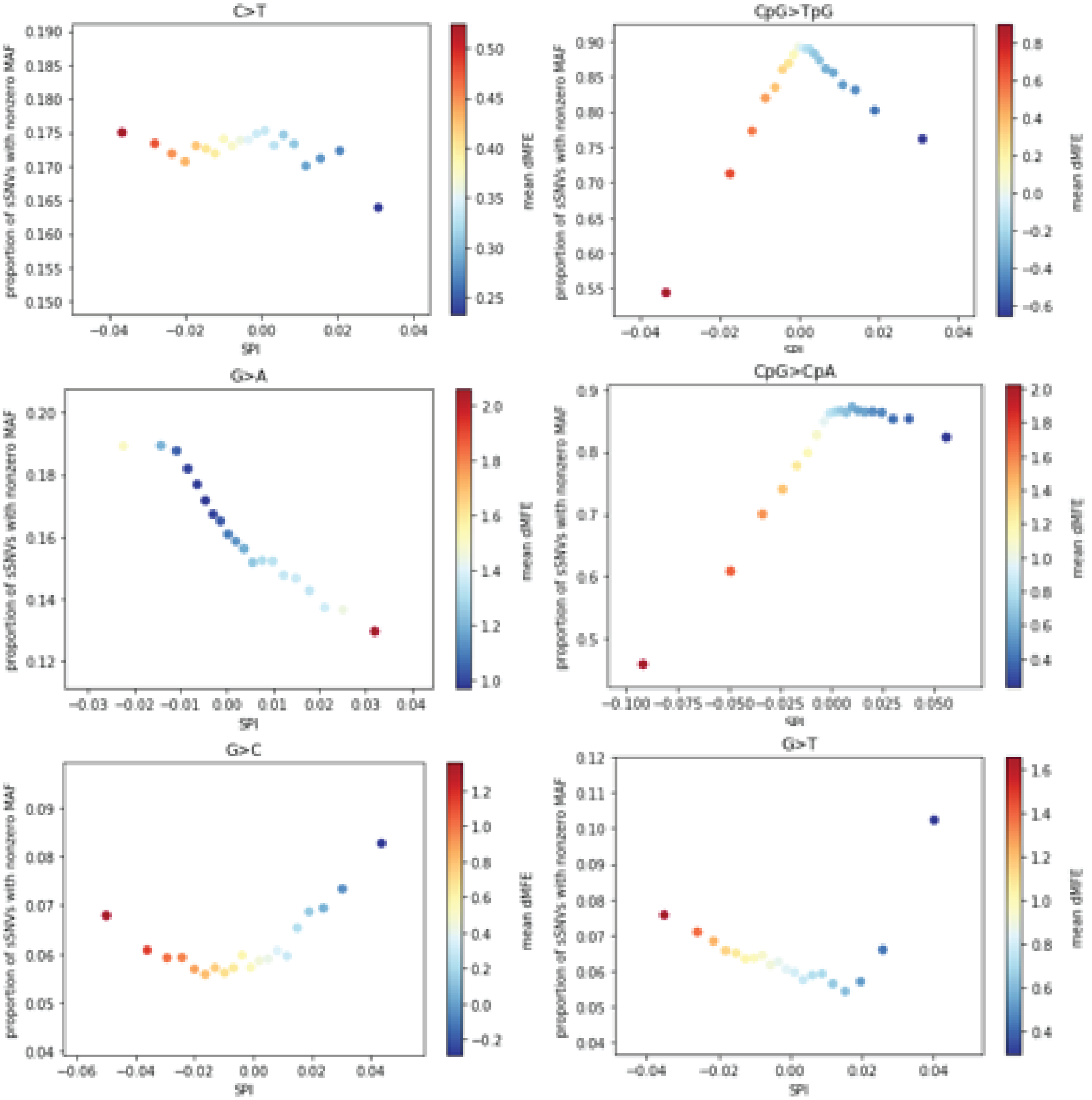

**Supplementary Data Figure 6. SPI score vs. P(Y=1).** In each of the 14 contexts from Table 1, we divide the set of sSNVs into 20 bins based on SPI score and then plot P(Y=1) vs. mean SPI score over each bin, coloring by dMFE.
